# The Host miR-17-92 Cluster Negatively Regulates Mouse Mammary Tumor Virus (MMTV) Replication via miR-92a by Targeting the Viral Genome

**DOI:** 10.1101/2023.08.31.555661

**Authors:** Jasmin Baby, Bushra Gull, Waqar Ahmad, Thanumol Abdul Khader, Neena G. Panicker, Shaima Akhlaq, Tahir A. Rizvi, Farah Mustafa

## Abstract

The mouse mammary tumor virus (MMTV) induces mammary tumors in mice. Recently, we have shown that MMTV perturbs expression of the host miR-17-92 cluster in MMTV-infected mammary glands and MMTV-induced mammary tumors. Known as oncomiR-1, this cluster is often dysregulated in a number of cancers, especially breast cancer. Therefore, we investigated whether there was a functional interaction between MMTV and the cluster. Our results reveal that MMTV expression led to dysregulation of the cluster in both mammary epithelial HC11 and HEK293T cells with the expression of miR-92a cluster member being affected the most. Conversely, over-expression of the whole or partial cluster significantly repressed MMTV expression. Interestingly, overexpression of miR-92a alone resulted in MMTV repression to the same extent as observed with the whole or partial cluster. Inhibition of miR-92a led to nearly a complete recovery of MMTV gene expression. Deletion/substitution of the miR-92a seed sequence within the cluster rescued MMTV expression. Dual luciferase assays identified MMTV genomic RNA as the target of miR-92a. These results show that the miR-17-92 cluster acts as part of the cell’s well-known miRNA-based anti-viral response to thwart incoming MMTV infection. Interestingly, miR-92a expression was the least amongst cluster members in MMTV-induced breast tumors, suggesting that the elevated level of MMTV expression in tumors is due perhaps to MMTV subverting miR-92a overexpression, restricting its ability to control MMTV replication during tumorigenesis. Thus, our study provides the first evidence demonstrating the biological significance of host miRNAs in controlling MMTV replication and influencing the process of tumorigenesis.

**Importance:** MMTV is a non-acute, slow-transforming retrovirus that causes breast cancer and lymphomas in mice. Consequently, understanding how MMTV interacts with its host can reveal critical aspects of cellular anti-viral responses and the process of tumorigenesis. This study reveals that the host miR-17-92 cluster plays an anti-viral role during MMTV infection which seems to be counteracted by MMTV during tumorigenesis. This observation is significant since MMTV-induced tumors share similarity with human breast cancer and the MMTV/mouse model is the best mammalian animal model to study breast cancer initiation, progression, and therapy. Moreover, MMTV is increasingly (though controversial) being implicated in human breast cancer, leukemias/lymphomas, and biliary cirrhosis due to its potential zoonosis into humans. Thus, if proven true, understanding how miRNAs modulate MMTV replication will be critical in appreciating how hosts miRNAs affect virus replication/tumorigenesis, and towards the development of novel miRNA-based therapeutics (e.g., anti-miRs) to thwart MMTV replication in humans.

## Introduction

RNA viruses have acquired various evolutionary advantages, the most important of which is their capacity to commandeer cellular machinery and elude host immune responses. These viruses are among the most common culprits responsible for clinically significant viral infections in humans, including influenza, human immunodeficiency virus (HIV), hepatitis B and C viruses (HBV & HCV), Ebola, Zika, Nipah, and more recently, the coronaviruses (SARS, MERS, COVID-19). Retroviruses are an exceptional class of RNA viruses with the capability to retro-transcribe their genomic RNA into double-stranded DNA, employing reverse transcriptase, an RNA-dependent DNA polymerase (Le Grice, 2012). The mouse mammary tumor virus (MMTV) is a *betaretrovirus* shown to cause breast cancer and lymphoma/leukemia in mice. Since its discovery in the 1930s, its antigens and sequences have been observed in human breast cancer as well, leading to the controversial possibility of a human mammary tumor virus (HMTV) (Amarante et al., 2019; Bevilacqua, 2022; Parisi et al., 2022). MMTV is a classic example of a slow-transforming retrovirus that causes cancer via insertional mutagenesis in mice over a period of 6-9 months where it integrates upstream of host genes involved in regulating growth and proliferation, such as *Wnt1, Fgf3, Rspo, and Notch4* (Dudley et al., 2016; Ross, 2010). Additionally, at least two structural genes of MMTV, *gag* and *env*, have also been reported to have oncogenic potential (Hook et al., 2000; Katz et al., 2005; Ross et al., 2006; Swanson et al., 2006). Despite extensive research into understanding MMTV replication since it’s discovery as a ‘milk-borne agent’ in 1936, there continues to be inadequacies in understanding the dynamics of viral replication, virus-host interactions, and pathogenesis, especially how MMTV interacts with the cellular RNA Interference (RNAi) machinery.

MicroRNAs (miRNAs) belong to the RNAi machinery, a new class of small non-coding RNAs that are super-regulators of gene expression (Flynt & Lai, 2008; Nilsen, 2007; Rani & Sengar, 2022). They silence gene expression by targeting mRNAs in a sequence-specific manner. Encoded within the genome, the biogenesis of miRNAs begins in the nucleus and closely follows the typical transcription activation and processing steps of any protein-coding mRNA, except that miRNAs additionally use a unique set of enzymes and proteins for this purpose (reviewed in Ha & Kim, 2014; O’Brien et al., 2018). The primary transcript normally produced by RNA polymerase II undergoes cleavage and is processed into a shorter 60-80nt precursor form by the nuclear RNase III enzyme, Drosha coupled with DGCR8, a double strand (ds) RNA binding protein. The precursor (pre-miRNA) form is shuttled into the cytosol by the export protein, Exportin-5, with the help of RAN-GTPase, another dsRNA binding protein. Once in the cytoplasm, the pre-miRNAs are cleaved by the cytoplasmic RNase III enzyme Dicer by a recognition element at the 3’ end of the pre-miRNA hairpin-loop (Koscianska et al, 2011). This process gives rise to the final dsRNA effector molecule, the mature miRNA, that comprises of a guide and a passenger strand. The guide strand forms a complex with the RNA-induced silencing complex (RISC), and identifies target mRNAs based on complementarity, which are then routed to cytoplasmic processing bodies (p-bodies) and marked for degradation. The two strands of the mature miRNA have distinct “seed” sequences and either of these have the potential to be loaded onto RISC to serve as the guide strand. This is an aspect critical in mRNA targeting and gene regulation, areas only now being unraveled (Medley et al., 2021).

Functioning as a part of the natural cellular RNAi machinery, miRNAs not only regulate changes in gene expression during development, but also serve as an “anti-viral” defense mechanism against invading viruses, a process that has been observed in a variety of taxa, including plants, nematodes, and arthropods (Czech et al., 2008; Ding & Voinnet, 2007; Hamilton & Baulcombe, 1999). The “anti-viral” miRNAs are induced in the presence of exogenous RNA in the host, which is identified by miRNA-viral RNA sequence complementarity in which the viral RNA is subjected to degradation by cellular enzymes (Lindsay, 2008; Toledo-Arana et al., 2007). The anti-viral potential of the RNAi machinery was first demonstrated in *C. elegans* where an inhibition of the vesicular stomatitis virus (VSV) was observed in mutant worms with an enhanced small RNA interference potential (Wilkins et al., 2005). Data from an HIV-1 study revealed that components of the RNAi machinery, Dicer and Drosha, suppressed viral replication, indicating that RNAi serves an anti-viral role in mammals (Triboulet et al., 2007). Viruses, on the other hand, can manipulate host miRNAs to prevent the cell’s anti-viral response or favor virus propagation (Frasca et al., 2022; Girardi et al., 2018; Grassmann & Jeang, 2008; Nahand et al., 2020). For example, HCV can significantly enhance levels of miR-373 in hepatocytes, which leads to attenuation of the JAK/STAT pathway via JAK1 and IRF9 at the RNA level, thereby counteracting interferon-mediated anti-viral responses in the cell (Mukherjee et al., 2015). Similarly, the human respiratory syncytial virus (RSV)-encoded proteins NS1 and NS2 can up-regulate cellular miR-29a to regulate the suppression of the interferon receptor, thus aiding viral replication stimulated by NS1 (Zhang et al., 2016).

While miRNAs are expressed from single genes, nearly 50% of miRNAs in *Drosophila melanogaster* and more than one-third of human miRNA genes are clustered together and expressed in polycistronic miRNA clusters, where a single parent transcript produces multiple mature miRNAs (Concepcion et al., 2012; Zhu et al., 2014; Tanzer & Stadler, 2004; Vilimova & Pfeffer, 2022). miR-17-92 is the first oncogenic miRNA cluster (oncomiR-1) to be described and studied extensively for its role in cell cycle regulation, differentiation, apoptosis, and tumorigenesis (Concepcion et al., 2012; de Pontual et al., 2011; Fang et al., 2017; Koralov et al., 2008; Mestdagh et al., 2010; Yang et al., 2016). Amplified in hematopoietic malignancies and solid tumors, a study of B-cell lymphomas in an Eµ-*myc* transgenic mice was the first to point out the cluster and its oncogenic potential (Ota et al., 2004; He et al., 2005). The miR-17-92 host gene (MIR17HG) is located in the non-transcribed region of human chromosome 13 (13q31.3) and chromosome 14 in mice (14; 14E4). The cluster is a polycistron, coding for a single RNA transcript that is processed into six mature miRNAs: miR-17, miR-18a, miR-19a, miR-20a, miR-19b-1, and miR-92a-1 (Ota et al., 2004). These mature miRNAs are conserved throughout vertebrates (Shi et al., 2013) and paralogous of the cluster exist, namely miR-106b-25 (7q22.1) and miR-106a-363 (Xq26.2). As mentioned earlier, miRNAs identify their target genes and exert post-transcriptional modulation via their seed sequences; i.e., nucleotides 2-8 of the mature form, to engage in the mRNA recognition/targeting event (Lewis et al., 2005). Based on their seed sequences, the miR-17-92 gene cluster and its homologs are categorized into four families: the miR-17 family (miR-17, miR-20a/miR-20b, miR-106a/miR-106b, and miR-93), the miR-18 family (miR-18a/miR-18b), the miR-19 family (miR-19a/miR-19b), and the miR-25 family (miR-25, miR-92a, and miR-363) (Mogilyansky & Rigoutsos, 2013).

With extensive research on the miR-17-92 cluster and its relevance as oncomiR-1 in tumor development and progression, in comparison, very little has been explored on the functional importance of this miRNA cluster in thwarting or facilitating viral infections. We have recently shown that the MMTV genome does not encode miRNAs; rather it dysregulates expression of host miRNAs in not only MMTV-infected lactating mammary glands, but also MMTV-induced tumors, including the miR-17-92 cluster (Kincaid et al., 2018). The current study was undertaken to study any functional interaction between MMTV and the miR-17-92 cluster at the cellular level and its consequences for virus replication. Since an upregulation of the cluster expression was observed in not only mouse mammary tumors that express one of the highest levels of the virus, but also infected lactating mammary glands (Kincaid et al., 2018), our hypothesis was that the cluster facilitates MMTV replication in mammary epithelial cells. However, we observed the opposite and found an anti-viral association between miR-17-92 and MMTV in infected cells, with the cluster leading to repression of MMTV expression. Furthermore, we identify miR-92a as one of the critical anti-viral components of the cluster that inhibits MMTV gene expression and replication by targeting its genomic RNA.

## Results

### The miR-17-92 cluster members are dysregulated upon MMTV expression

To explore whether MMTV could interfere with the expression of the miR-17-92 cluster, we looked into our miRNAseq data of a mouse mammary epithelial cell line HC11 compared to the same cell line expressing MMTV, HC11-MMTV (Baby, 2022; Gull et al., manuscript submitted). Mammary epithelial cells are the most permissive cells for MMTV infection *in vivo,* responsible for ensuring the passage of the virus from the mother to the pups via breast milk; additionally, these are the main cells targeted for mammary tumorigenesis in mice (Dudley et al., 2016; Ross, 2010). Figure 1A reveals the heat map of the expression of the cluster members in the two cell lines (two biological replicates, 1 & 2, are shown for each). As can be seen, a distinct dysregulation of the cluster member expression could be observed upon MMTV expression. Fold-change analysis of differential gene expression between the two cell lines revealed that while the expression of some of the cluster members was not much affected (miR-17-3p and miR19a-3p), a distinct up-regulation could be observed in the expression of other cluster members, such as miR-18a and miR-92a, with miR-92a showing the highest induction at 4-folds upon MMTV expression (Fig. 1B). Other cluster members not shown in Fig. 1B, did not demonstrate any substantial levels of expression in the HC11 cells.

**Figure 1:**
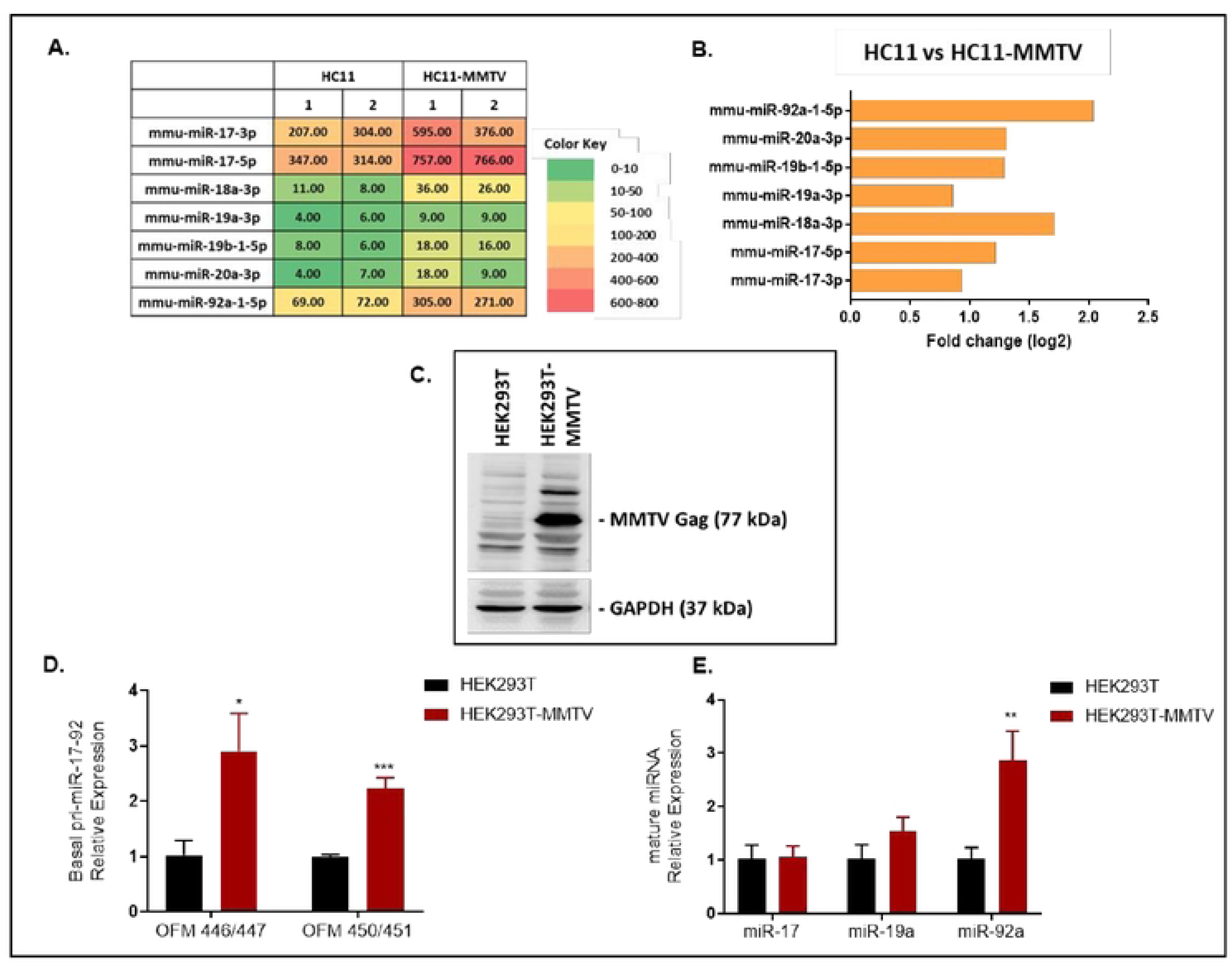
The miR-17-92 cluster members are dysregulated upon MMTV expression. **(A)** Heat-map of miR-17-92 cluster upon small miRNAseq analysis of the normal mouse mammary epithelial cell lines, HC11 and HC11-MMTV. The miRNAseq was conducted on two biological replicates shown as 1 and 2. The numbers in the table represent the normalized reads observed upon miRNAseq. **(B)** Fold-change analysis of mature miR-17-92 cluster member expression using the miRNAseq data from HC11 cells. The Q-value (equivalent of *p*-value corrected for multiple test hypothesis) for each of the miRNA shown was statistically significant (Q<0.005). **(C)** Western blot analysis of HEK293 and HEK-293T cells expressing MMTV using an anti-MMTV Gag antibody. An antibody against a housekeeping gene (GAPDH) was used as a loading control. **(D)** Quantitative RT-PCRs (RT-qPCRs) to assess endogenous levels of the primary form of miR-17-92 cluster upon MMTV expression in HEK-293T cells. β-Actin was used as the endogenous control. **(E)** RT-qPCR analysis of mature miR-17, miR-19a and miR-92a in HEK293T cells. U6 was used as the endogenous control. All experiments were carried out in triplicates. Statistical significance is shown as * where *p is ≤ 0.05; **p ≤ 0.01, and ***p ≤0.001.

Being of mouse and mammary epithelial origin, HC11 are the most appropriate cells to study MMTV replication in its relevant context; however, they express endogenous strains of MMTVs that are found ubiquitously in most mouse species (Dudley et al., 2016; Ross, 2010). These endogenous strains of MMTV (specifically *mtv-6, mtv-8,* and *mtv-9* in HC11 since they are of BALB/c origin) are defective for replication, but are able to express parts of their genomes that could have confounded our results (Holt et al., 2013; Salmons et al., 1986). Therefore, to study the interaction between MMTV and the miR-17-92 cluster further, we decided to use the human embryonic kidney cell line HEK293T as being of human origin, it is devoid of any endogenous strains of MMTV.

Towards this end, similar to HC11, the HEK293T cells were transfected with HYBMTV, a replication-competent molecular clone of MMTV (Shackleford & Varmus, 1988), to establish an HEK293T-MMTV stable cell line constitutively expressing the virus (Fig. 1C). Quantitative real-time PCR with two primer sets, primer pair 1 (OFM 446/447) and primer pair 2 (OFM450/451), that targeted the primary (pri)-form of the cluster, revealed that MMTV significantly enhanced expression of the primary form of the miR-17-92 cluster by more than 2.5-fold relative to HEK293T cells alone (Fig. 1D). Subsequently, the levels of some of the mature cluster members (miR-17, miR-19a and miR-92a) were assessed using TaqMan miRNA assays to determine if the level of mature miRNAs followed the same course. Similar to the HC11 cells, a significant 2.9-fold increase in the endogenous levels of miR-92a was observed, while levels of miR-17 and miR-19a were not affected significantly (Fig. 1E). These data reveal that MMTV enhanced expression of the primary form of the miR-17-92 cluster, while its effect on individual mature cluster members varied with miR-92 being affected the most. They also confirm that observations made in the HEK293T cells recapitulated what was observed in the more relevant HC11 cells, making our HEK293T system valid for further investigation of the interactions between MMTV and the miR-17-92 cluster members.

### MMTV expression is significantly down-regulated upon miR-17-92 over-expression

Previously, it has been shown that host miRNAs can have anti-viral potential where activation of these miRNAs leads to reduction in virus expression (Fu et al., 2019; Jung et al., 2013; Mallick et al., 2009; Triboulet et al., 2007). Since miR-92a was observed to be the most affected cluster member upon MMTV expression in both cell lines, we asked whether miR-92a could have a negative feedback relationship with viral replication as an anti-viral miRNA. To test this hypothesis, expression plasmids carrying either the full cluster, a truncated version of the cluster expressing miR-19a-20a-19b (miR-19/20) but lacking miR-92a, or miR-92a only were transfected into HEK293T-MMTV cells to create miRNA over-expression (OE) stable cell lines along with a control expressing the empty vector (EV) without any miRNA insert (Fig. 2A & 2B). The miRNA over-expression phenotype was confirmed using TaqMan miRNA assays and its effect on MMTV expression was assessed at the RNA and protein levels.

**Figure 2:**
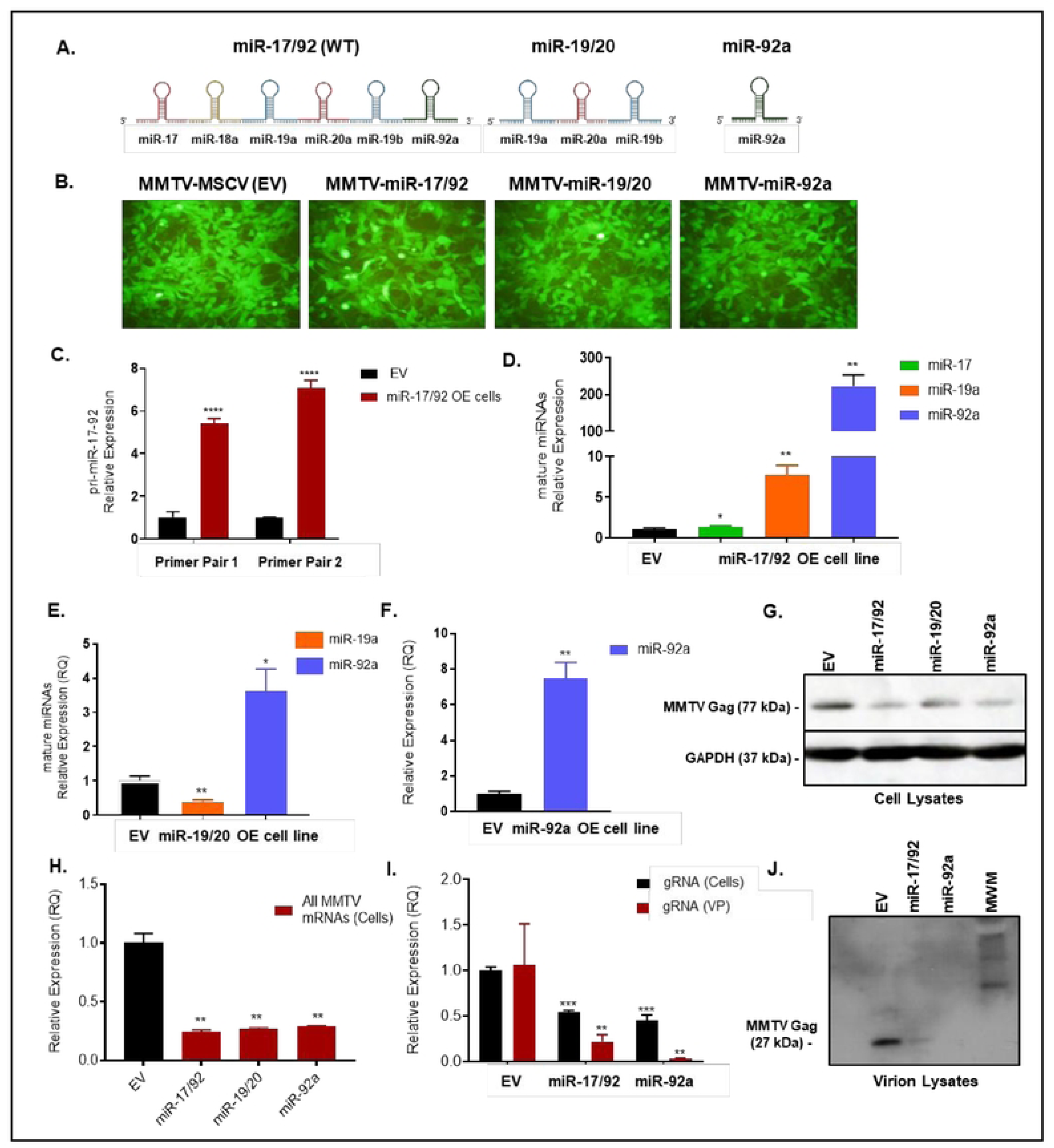
Over-expression of the miR-17-92 cluster down-regulates MMTV expression. **(A)** An Illustration of the exogenously-expressed miR-17-92 cluster, its truncated form miR-19a-20a-19b, and miR-92a over-expressed (OE) in the HEK293T-MTMV cells. **(B)** GFP-positive stably-transfected cells selected in puromycin and visualized using a fluorescent microscope (10x). **(C)** Expression of pri-miR-17-92 in miR-17-92 OE cell line quantified using a SYBR Green qPCR assay. **(D)** TaqMan miRNA assays used to quantify expression of mature miR-17, miR-19a, miR-92a in the miR17-92 OE cell line, **(E)** miR-19/20 OE cell line, and **(F)** miR-92a OE cell line. U6 snRNA was used as the endogenous control in these assays. **(G)** Western blot analysis of MMTV Gag expression in cell lysates and **(J)** viral particles extracted from the respective miRNA OE stable cell lines. GAPDH was used as the endogenous control. **(H)** MMTV RNA-specific TaqMan assay to quantify all MMTV messages in the three OE cell lines. **(I)** MMTV genomic RNA was assessed using gRNA-specific MMTV TaqMan assay in the miR-17-92 cluster OE and the miR-92a OE cell lines. β-Actin was used as the internal control in the TaqMan assays. The empty vector (EV) was designated as 1 in all analyses. Mean ± SD (n=3). Statistical significance is shown as * where *p is ≤ 0.05; **p ≤ 0.01, ***p ≤0.001, ****p≤ 0.0001.

As can be seen, the miR-17-92 full cluster OE cells showed a significant increase in the amounts of pri-miR-17-92 produced (5-7 folds) compared to the EV-expressing cells (Fig. 2C). Next, we analyzed the expression of some of the individual members of the cluster in this cell line. A statistically significant over-expression of cluster members miR-17 and miR-19a could be observed in the 1.3- to 7.7-fold range; however, miR-92a was upregulated the most (~225-folds) (Fig. 2D). The miR-19/20 cell line expressing the partial cluster (miR-19a-20a-19b), showed an unexpected down-regulation in the levels of miR-19a (Fig. 2E), while the miR-92a over-expression cell line confirmed a clear and significant 7-fold over-expression of miR-92a (Fig. 2F). Next, the functional consequence of miR-17-92 cluster over-expression on MMTV expression was assessed at the protein and RNA levels. Test of the MMTV Gag structural gene expression levels were observed to be down-regulated at the protein level when compared to the empty vector control in all the three OE cell lines (Fig. 2G). Densitometric analysis on the same revealed that the level of repression of MMTV expression by miR-92a alone was comparable to that of its expression from the full cluster (data not shown). This down-regulation of MMTV expression was confirmed at the RNA level where over-expression of either the full miR-17-92 cluster, the miR-19/20 partial cluster, or miR-92a only significantly down-regulated MMTV mRNA expression (Fig. 2H). Subsequently, the effect of the full cluster and miR-92 over-expression on MMTV virus particle production and genomic RNA packaging was assessed using RT-qPCR. As can be seen, the amount of genomic RNA expressed in the cells was significantly down-regulated in both the miR-17-92 full cluster and miR-92a over-expressing cells. This decrease was reflected in reduced amounts of gRNA packaged into the virus particles with the miR-92a over-expressing cells showing the most inhibition (Fig. 2I). This observation was confirmed by western blot analysis of virus particles produced in each cell line. As can be seen, the level of virions produced in the miR-17-92 cluster and miR-92a over-expressing cells was drastically reduced compared to the control EV-expressing cells (Fig. 2J). Considering that the unspliced, full-length genomic RNA in retroviruses is used not only to produce the structural polyprotein such as Gag, but also for its encapsidation into the newly-assembling virus particles, this result suggests that expression of the miR-17-92 cluster, and in particular its miR-92a component, results in down-regulation of virus replication, perhaps as an anti-viral response of the cells to MMTV infection.

### Anti-miRNA oligo-based miR-17-92 and miR-92a inhibition rescues MMTV expression

To validate these findings, we took the reverse approach where modified anti-miRNA oligos were used to inhibit mature forms of cluster members in the three different OE cell lines. A combination of anti-miR-17, -19a and -92a inhibitors were used at 15 pmol concentration to inhibit miRNA abundance in the miR-17-92 HEK293T-MMTV OE cell line. Similarly, 15 pmol of anti-miR-19a and 15 pmol of anti-miR-92a were used individually to suppress miRNA overexpression in the miR-19/20 and miR-92a OE cell lines, respectively. TaqMan miRNA assays were carried out to quantify and confirm the inhibition of miR-17, miR-19a and miR-92a. Interestingly, in the miR-17-92 OE cell line treated with the anti-miRNA cocktail, no significant reduction in targeted miRNAs was observed (Fig. 3A), while, the miR-19/20 OE cell line treated with anti-miR 19a showed a downward trend of miR-19a suppression that was statistically significant (Fig. 3B). The most significant inhibition, however, was observed in the miR-92a OE cell line treated with anti-miR-92a where ~80% decrease in miR-92a population was observed (Fig. 3C). Despite variable levels of respective miRNA inhibition obtained, we could discern some rescue of MMTV Gag (Pr77) expression in all the cell lines treated with the respective anti-miR oligo inhibitors compared to the scramble negative control (NC1-3) at the protein level (Fig. 3D).

**Figure 3:**
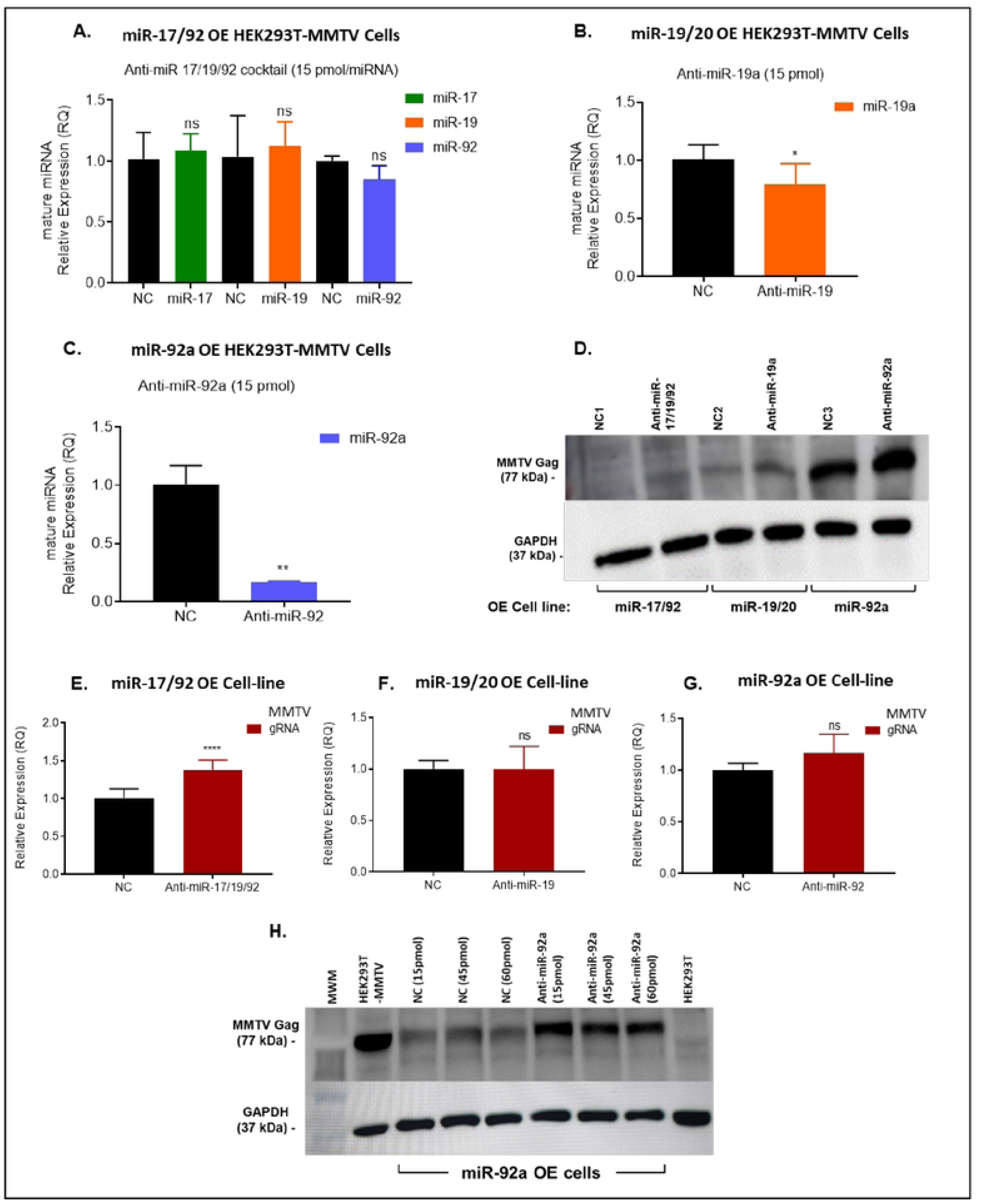
miR-17-92 cluster and miR-92a inhibition rescues MMTV genomic RNA expression. TaqMan miRNA assays were used to quantify: **(A)** miR-17, miR-19a and miR-92a in miR-17-92 over expression (OE) cell line, **(B)** miR-19a in the miR-19a-20a-19b OE cell line, and **(C)** miR-92a in the miR-92a OE cell line. U6 snRNA was used as the endogenous control in the miRNA RT-qPCR assays. The error bars represent ± SD. **(D)** Western blot analysis of whole cell lysates (40 µg) from the miRNA OE cell lines treated with miRNA inhibitors for MMTV Gag expression. GAPDH was used as the internal control. Expression of the MMTV genomic RNA was quantified in the: **(E)** miR-17-92 OE cells treated with a cocktail of anti-miR-17, -19, & -92a oligos, **(F)** miR-19a-20a-19b OE cells treated with anti-miR-19a oligos, and **(G)** miR-92a OE cells treated with anti-miR-92a oligos. β-Actin was used as the internal control in the MMTV TaqMan RT-qPCR assays. The empty vector (EV) was designated as 1 in all analyses. Mean ± SD (n=3). **(H)** Western Blot analysis of whole cell lysates (50 µg) from the miR-92a OE cell line treated with anti-miR-92a oligos at varying concentrations. GAPDH was used as the internal control. Statistical significance is shown as * where *p is ≤ 0.05; **p ≤ 0.01, ***p ≤0.001, ****p≤ 0.0001. ns, not significant; ns (P > 0.05).

To analyze the effect of the anti-miR oligo inhibitors on virus expression at the transcriptional level, total RNA from the three cell lines was subjected to MMTV-specific genomic RNA RT-qPCR. As can see seen, a significant up-regulation in MMTV gRNA expression was observed in the miR-17-92 OE cell line treated with anti-miR-17+ anti-miR-19a + anti-miR-92a inhibitor cocktail compared to the negative scramble control treatment (Fig. 3E). However, no significant rescue of MMTV gRNA expression could be observed with treatment of the miR-19/20 OE cells with the anti-miR-19 oligo (Fig. 3F). In contrast, some rescue could be observed of MMTV gRNA expression with the anti-miR-92a inhibitor in the miR-92a OE cell line, though it was not statistically significant (Fig. 3G). To determine whether the effect of anti-miR-92a was real, the experiment was repeated with increasing doses of the inhibitor oligo in the miR-92a OE cells. As anticipated, a significant rescue of MMTV Gag protein expression could be observed upon increasing amounts of the anti-miR-92a inhibitor oligo that could be visualized on a western blot (Fig. 3H). Although not dose-dependent, these observations confirm that the miR-17-92 cluster suppresses MMTV gene expression via miR-92a.

### Plasmid-based miRNA inhibition (PMIS) of miR-17-92 cluster members up-regulates MMTV expression

To investigate these findings further, a plasmid-based approach to miRNA inhibition was used in which vectors expressing anti-sense miRNAs were used that repress miRNAs functionality by forming a stable PMIS-miRNA complex (Cao et al., 2016). Towards this end, three stable PMIS cell lines were established in HEK293T-MMTV cells against miR-17 (PMIS 17), miR-92a (PMIS 92) and empty control vector (EV) that constitutively expressed the anti-sense miRNA or the empty vector as a control. These cell lines were tested for the expression of the respective miRNAs inhibited using specific TaqMan miRNA assays (Fig. 4). Interestingly, these assays revealed that the PMIS vectors could not down-regulate the targeted endogenous miRNA population; i.e., while a slight downward trend could be observed of miR-17 that was not statistically significant, an up-regulation of ~1.5-folds was observed for miR-92a which was statistically significant (Fig. 4A & 4B). This was surprising and we wanted to test whether this observation was real or an artefact of the PMIS vector system. We hypothesized that the miRNA-anti-sense PMIS complex probably forms a duplex that is stable and was being detected in these assays. Therefore, despite a physical lack of inhibition of the test miRNAs, miR-17 and miR-92a, we tested for a functional loss of the tested miRNAs. This was achieved by analyzing the expression of the validated target of miR-17, the phosphatase and tensin homolog (PTEN) mRNA. As can be seen, a 2-fold up-regulation of PTEN transcript was observed in the PMIS-17 cell line that was statistically significant, confirming the functional inhibition of miR-17 in the PMIS stable cell line (Fig. 4C). Based on these results, we analyzed MMTV expression in PMIS-17 and PMIS-92 stable cell lines. As expected, despite a lack of physical down-regulation of the targeted miRNAs, we observed an up-regulation of the MMTV Gag protein and gRNA in a statistically significant manner (Fig. 4D & 4E). These results confirm the suppressive role of miR-17 and miR-92a in MMTV gene expression.

**Figure 4:**
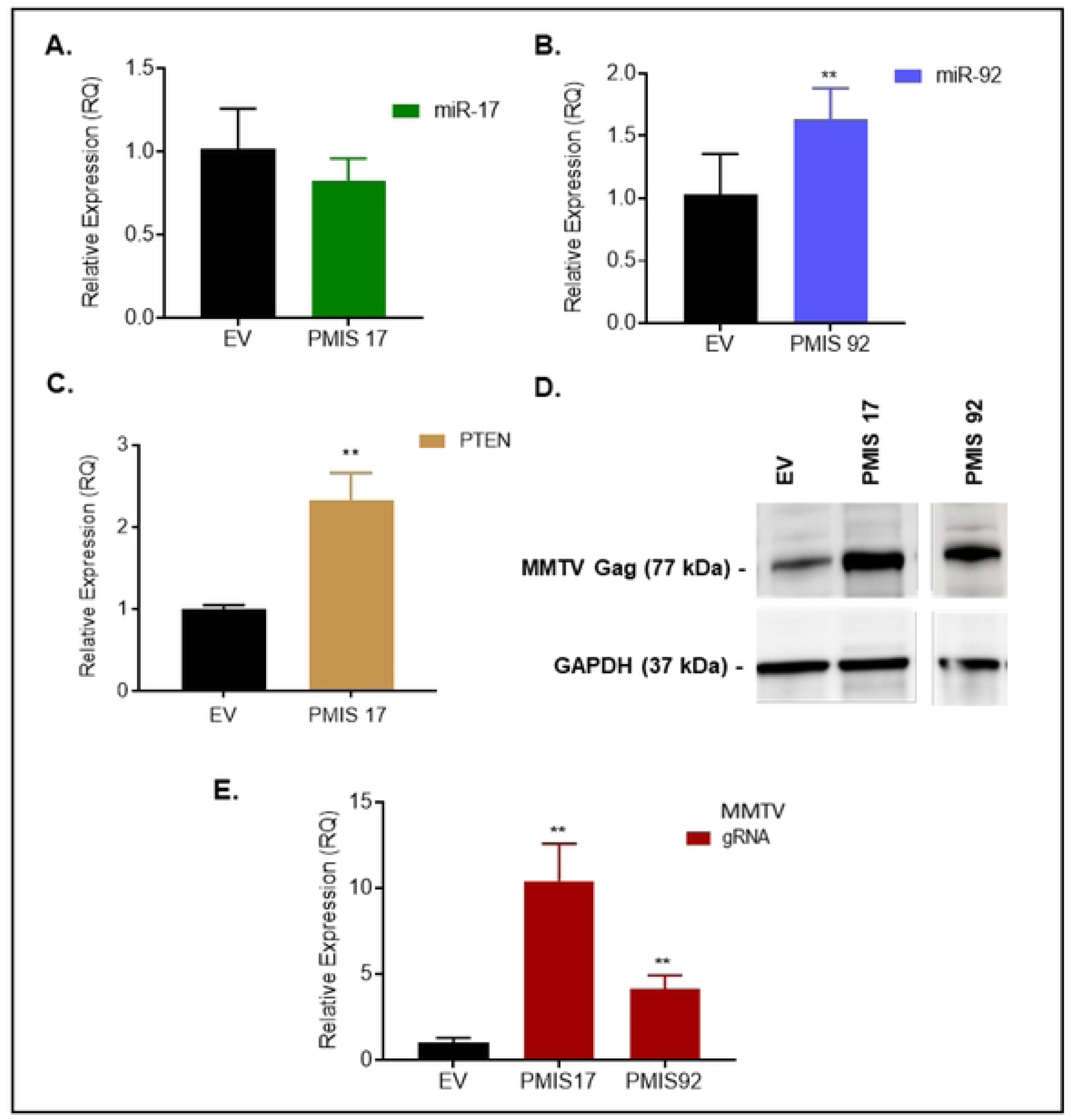
Rescue of MMTV expression upon plasmid-based miRNA inhibition. **(A)** Mature miR-17 and **(B)** miR-92a levels were quantified in their respective PMIS inhibitor expressing cell lines using TaqMan miRNA qPCRs. U6 snRNA was used as the endogenous control. **(C)** Expression of miR-17 target, PTEN was quantified using RT-qPCR; β-Actin was used as the internal control. **(D)** Western blot analysis of MMTV Gag expression across the three cell lines using 50 µg protein. GAPDH served as the endogenous control. **(E)** The expression of all MMTV transcripts were quantified using RT-qPCRs with β-Actin as the internal control. The empty vector (EV) was designated as unit 1 in all experiments with means represented as ± SD (n=3). Statistical significance is shown as * * where *p is ≤ 0.05 and **p ≤ 0.01.

### miR-92a is a critical anti-viral component of the miR-17-92 cluster

Next, we used a mutational approach to study the role of miR-92a as an anti-viral cellular component against MMTV. Plasmid vectors carrying the wild-type miR-17-92 (WT) cluster, miR-17-19b (Δ92a)-a miR-92a-deleted version of the whole cluster, and a substitution mutant of the miR-92a seed sequence in the cluster, miR-17-92aMUT, were transiently transfected into HEK293T-MMTV cell line to determine whether they could affect MMTV expression. As expected, expression of the WT miR-17-92 cluster down-regulated MMTV expression at the protein level compared to the empty vector control (EV) (Fig. 5A). This phenotype was reversed when the miR-92 seed sequence was either deleted from the cluster or mutated via a substitution (Fig. 5B). Quantitation of MMTV expression by the TaqMan RT-qPCR assay confirmed these results where expression of the wild type cluster resulted in down-regulation of MMTV expression in a statistically-significant manner, while either deletion or mutation of the miR-92a component of the cluster rescued the levels to wild type levels (Fig. 5C). These results clearly demonstrate that miR-92a is the major anti-viral component of the miR-17-92 cluster in controlling MMTV replication in the cell.

**Figure 5:**
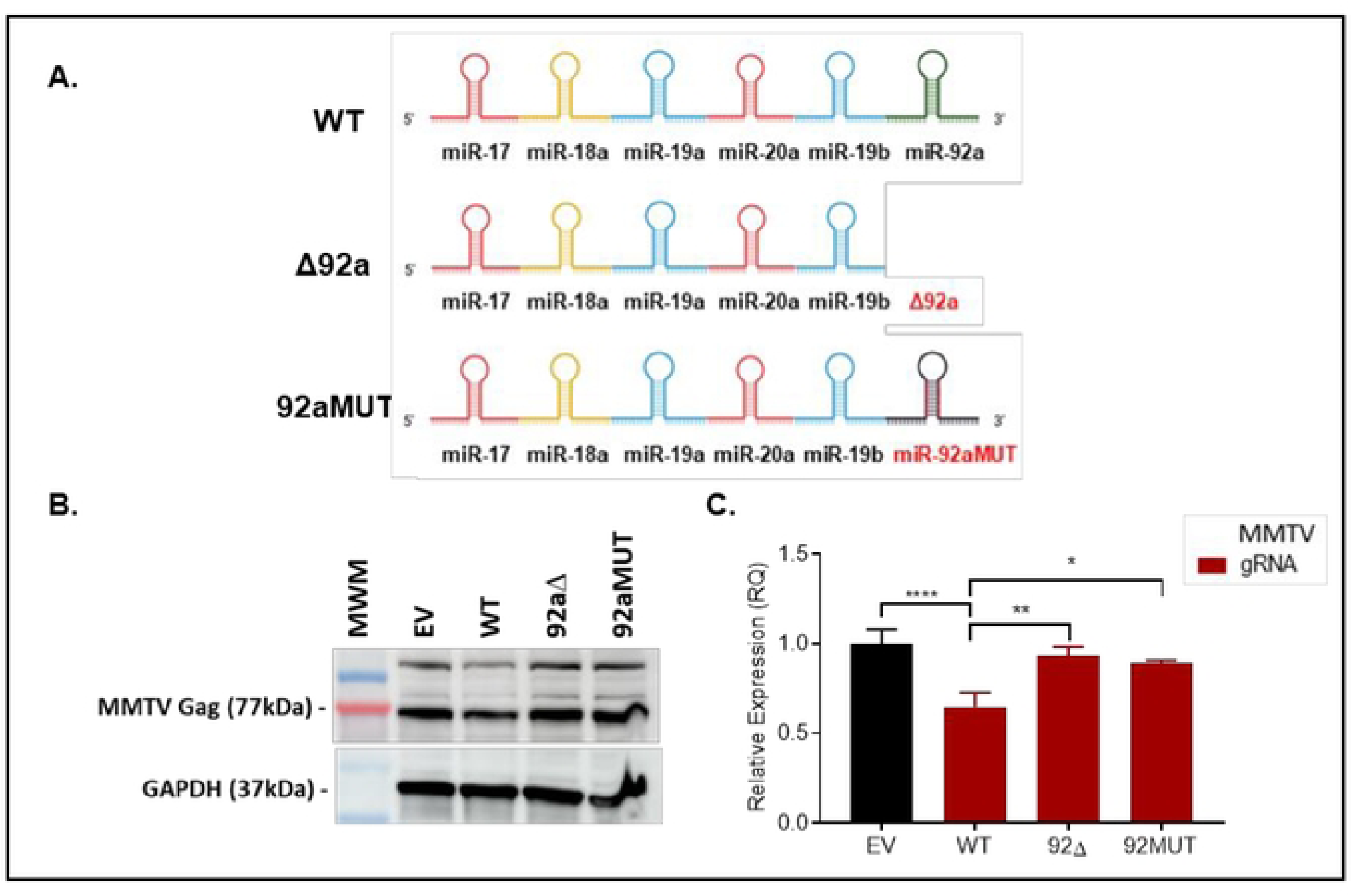
MMTV expression rescued upon deletion or mutation of miR-92a seed sequence. **(A)** Schematic representation of the full miR-17-92 cluster (WT), or cluster with deletion of miR-92a (Δ92a) highlighted in red, or a substitution mutation of miR-92a (92aMUT) within the cluster highlighted in red. **(B)** Expression of MMTV Gag quantified by western blot analysis using 40 µg lysate from the different transiently transfected cell lines; GAPDH was used as the internal control. **(C)** Quantification of all MMTV mRNAs by RT-qPCR. β-Actin used as the internal control. The empty vector (EV) was designated as unit 1 with means represented as ± SD (n=3). Statistical significance is shown as * where *p is ≤ 0.05; **p ≤ 0.01, ***p ≤0.001, and ****p≤ 0.0001.

### The miR-17-92 cluster directly interacts with the MMTV genome

Following the demonstration of a repressive role of miR-92a in MMTV expression, we aimed at identifying its probable target sites on the MMTV genome using Sfold STarMir, a crosslinking immunoprecipitation (CLIP)-based prediction software with combined algorithms of RNAhybrid and logistic thermodynamic probability (Rehmsmeier et al., 2004; Rennie et al., 2016). Bioinformatic analysis of the data revealed thousands of putative binding sites of the miR-17-92 cluster members on the viral genome. The application of stringent parameters (ΔGhybrid: ≥ −20kcal/mol; Logistic probability score above 0.7) reduced these to a total of 74 high probability binding sites that were spread across the MMTV genome and targeted all the known mRNAs of the virus, including the genomic RNA that also serves as the mRNA for the structural genes Gag/Pro/Pol (Fig. 6A). These predicted miR-17-92 cluster binding sites belonged to specific members of the miR cluster, including miR-17, miR-18a, miR-19b, miR-20a, and miR-92a, but excluded miR-19a (Fig. 6B). Interestingly, roughly two-thirds (62% or n=46) of all predicted binding sites belonged to miR-92a, while other members such as miR-17 and miR-19b had the least sites (Fig. 6B). The 46 miR-92a binding sites were observed to be spread throughout the MMTV genome and targeted different mRNAs of the virus. Of these 46 miR-92a binding sites, ten sites were observed within the Gag region of the MMTV genome; however, one particular binding site had the highest logistic and thermodynamic probability of being a functional binding site with an 8-mer nucleotide seed type (Fig. 6C), suggesting that the Gag region was a particular target of miR-92a. This region along with the regions encoding Pro and Pol are found exclusively within the full-length genomic RNA of the virus and is spliced out from all the other mRNAs of the viral genome (the blue boxed region in Fig. 6A). Thus, this suggested that rather than a particular mRNA of the virus being the target of these miRNAs, it was the genomic RNA that was being targeted by the cluster members. Figure 6B highlights all the miRNAs that targeted the Gag/Pro/Pol regions. As can be seen, 40 of the total 74 miRNA binding sites (54%) were present within the Gag/Pro/Pol region of which only Gag had 16 specific binding sites, 10 of which belonged to miR-92a, including the functional binding site shown in Fig. 6C.

**Figure 6:**
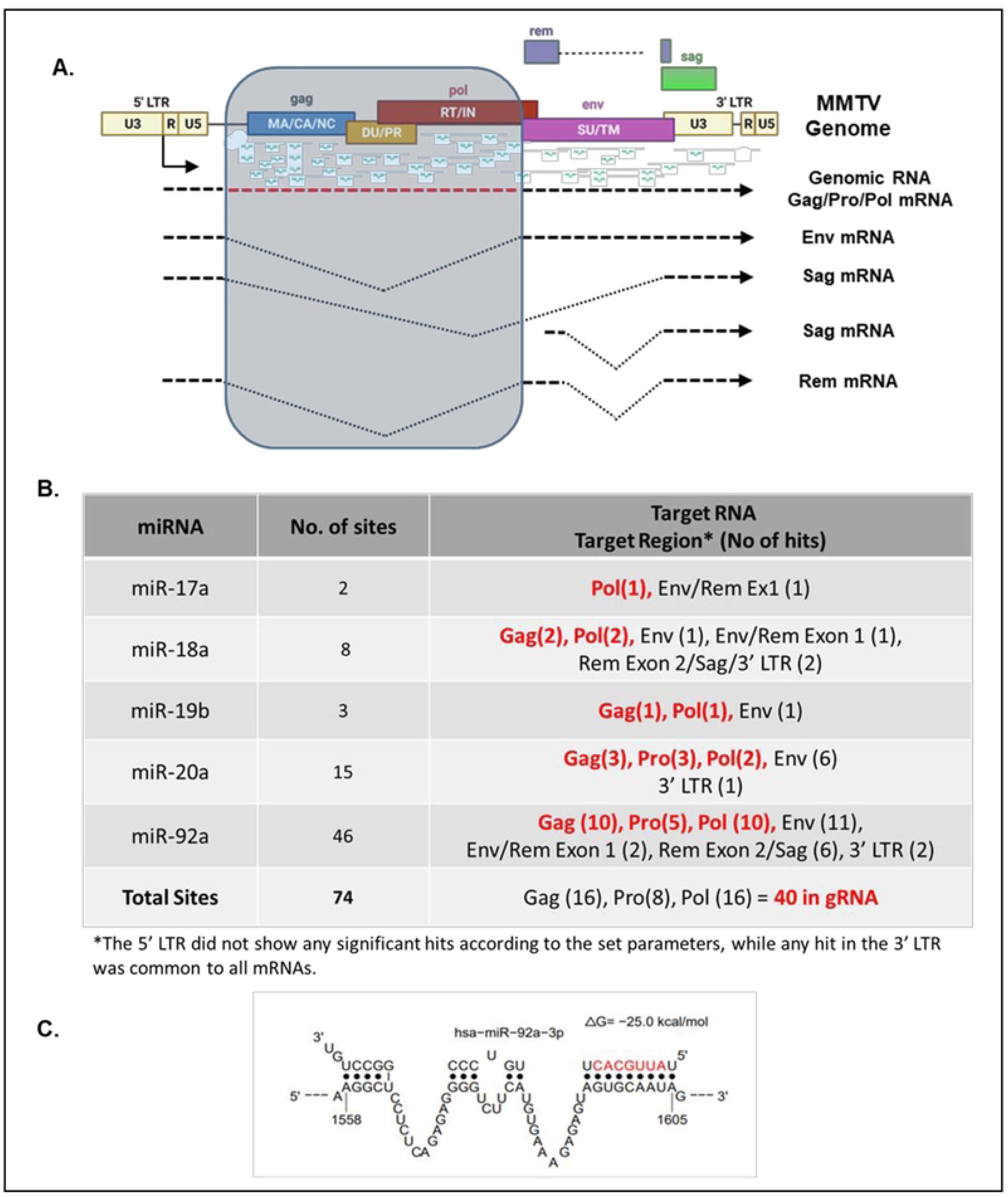
An *in silico* analysis of the potential of the miR-17-92 cluster members to target the MMTV genomic RNA using STarMir bioinformatic tool. (A) Illustration of the MMTV genome and its various mRNAs along with the potential miR-17-92 cluster member predicted binding sites. The shaded box in blue highlights the region of the MMTV genome exclusively found in the full-length genomic RNA which is the same as the mRNA for Gag/Pro/Pol viral proteins. **(B)** Characterization of the miRNAs predicted to target the MMTV mRNAs. The target RNA regions highlighted in red belong exclusively to the full-length Gag/Pro/Pol mRNA or the gRNA with the number of target sites observed in parenthesis. **(C)** Illustration of the highest probable binding site of miR-92a on MMTV Gag.

To test the validity of these predictions and explore the possibility of a direct interaction between miR-92a and the MMTV genome, the Gag region of the MMTV genome present exclusively in the full-length genomic RNA and containing the predicted functional binding site, in addition to the 15 others, was cloned into the miRNA target plasmid, pmirGLO (Promega), creating GagGLO (Fig. 7A). The Gag region was inserted in between the *firefly luciferase* gene and the SV40 poly A sequences, thus creating a fused Luciferase-Gag transcript containing the putative miR-92a target sites. The pmirGLO vector contains a second luciferase gene, *renilla luciferase*, expressed from an independent promoter that was used to internally normalize the transfection efficiencies. The GagGLO plasmid was tested in parallel with the control pmirGLO vector in the HEK293T and HEK293T-MMTV cells in transient transfection assays to determine whether presence of enhanced levels of cluster members in the HEK293T-MMTV cells would target the hybrid GagGLO mRNA in the MMTV-expressing cells, resulting in reduction in the luciferase activity compared to its expression in the HEK293T cells devoid of MMTV expression.

**Figure 7:**
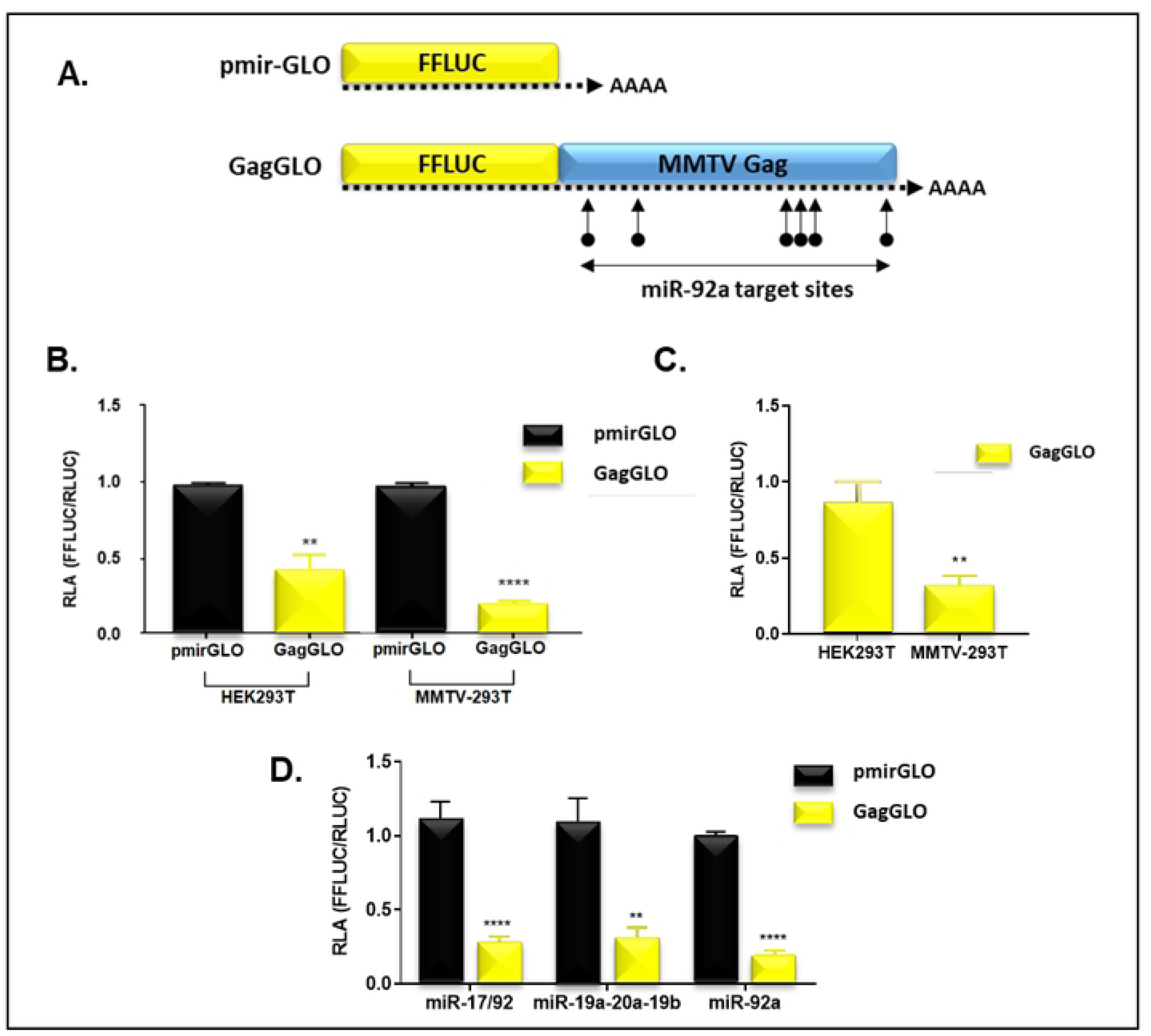
Dual-Luciferase activity-based interaction assays to test the direct interaction of the miR-17-92 cluster with the MMTV genome using. **(A)** Illustration of the GagGLO constructs and transcripts produced in the presence or absence of the gag *gene* cloned downstream of the *firefly luciferase* gene. The parent pmirGLO control vector also has an internal *renilla luciferase* gene cassette serving as transfection control for experiments. The over-expression reporter vectors, pmiRGLO (control) and GagGLO were transfected into **(B)** HEK293T, MMTV-293T and dual luciferase activity was determined 48 hours post-transfection. **(C)** A re-comparison of relative luciferase activity of GagGLO in HEK293T (control) and MMTV-293T cells shown in panel B. **(D)** Test of the pmiRGLO constructs in the three specific miRNA overexpressing (OE) cell lines described earlier. The empty vector, pmiRGLO (EV) was designated as unit 1 with means represented as ± SD (n=3). Statistical significance is shown as * where *p is ≤ 0.05; **p ≤ 0.01, ***p ≤0.001, ****p≤ 0.0001.

As can be seen, the levels of normalized firefly luciferase activity from GagGLO in the HEK293T-MMTV cells was indeed significantly lower compared to its expression in the normal HEK293T cells (Fig. 7B). Interestingly, even in the HEK293T cells devoid of MMTVs, there was a significant down-regulation of GagGLO, most likely due to the presence of endogenous miRNAs like miR-92a and others that may target the Gag region. The luciferase activity of GagGLO was reduced by 80% in the HEK293T-MMTV cells compared to HEK293T cells, pointing to the potential activation of miR-92a expression in the presence of MMTV (Fig. 7C). The same effect was observed when these two vectors were tested in the three cell lines that over-expressed either the whole miR-17-92 cluster, the truncated cluster (miR-19/20), or miR-92a only, thus mimicking MMTV infection which up-regulates miR-92a (Fig. 7D). These results clearly demonstrate that the Gag region of the MMTV genome, a region that is part of the full-length RNA and removed from the various spliced mRNAs of MMTV, is a direct target of the miR-17-92 cluster members, more specifically miR-92a.

## Discussion

The current study stemmed from our observations that MMTV disrupts cellular miRNA expression in the host (Kincaid et al., 2018). Both infected mammary glands and MMTV-induced tumors, tissues with high level MMTV expression, were observed to express high levels of several miR-17-92 cluster members, suggesting a possible interaction between the cluster and the virus that facilitated virus replication and presumably tumor induction. This encouraged us to explore the role of miR-17-92 cluster in MMTV replication with the hypothesis that the miR-17-92 cluster facilitates MMTV replication using functional assays that manipulated endogenous levels of miR-17-92 (Figs. 2-4) as well as mutational analysis (Fig. 5). However, rather than finding a “pro-viral” role, we identify an anti-viral role of the miR-17-92 cluster in MMTV replication where the cluster member, miR-92a, was observed to target the full-length MMTV genomic RNA, thereby drastically suppressing MMTV replication (Figs. 6 & 7). Thus, the miR-17-92 cluster seems to act as part of the cell’s anti-viral response, thwarting incoming virus infections for which miRNAs are well known for (Czech et al., 2008; Ding & Voinnet, 2007; Hamilton & Baulcombe, 1999).

While these observations explain how the miR-17-92 cluster functions an “anti-viral” response to suppress MMTV replication in cell lines, they do not explain our observation of high levels of MMTV expression in the presence of up-regulation of the miR-17-92 cluster members in both the infected mammary glands and mammary tumors of MMTV-infected mice (Kincaid et al., 2018). We speculate that upon initial MMTV infection, the cells mount an anti-viral response to suppress MMTV infection by activating the miR-17-92 cluster, and in particular miR-92a. However, the virus is able to overcome this restriction by subverting the expression of miR-92a, leading to enhanced virus expression that eventually leads to mammary tumorigenesis. This speculation suggests that the elevated levels of MMTV in mammary tumors, despite the up-regulation of cluster members, could be due to lower levels of miR-92a expression in the tumor samples.

To test this hypothesis, retrospection into the small RNAseq data from MMTV-induced tumors conducted earlier (Kincaid et al., 2018) revealed interesting expression patterns of the miR-17-92 cluster members which further gives confidence to our assertion. Intriguingly, while an increased expression of the oncogenic miR-19 family members, miR-19a and miR-19b, could be observed, the expression of miR-92a was the least amongst the expressed cluster members (Fig. 8). This suggests that the major anti-viral component of the miR-17-92 cluster was suppressed in MMTV-induced mammary tumors compared to uninfected mammary glands. A similar observation has been frequently demonstrated in different sub-types of breast cancers, with members miR-17, miR-19 and miR-20 implicated in tumor induction and progression (Moi et al., 2019). The differential expression of the various members of a miRNA cluster is tightly regulated in a cell, cluster, and context-dependent manner where the type of trans factors available within a cell (proteins & long non-coding RNAs), the diseased state of the cell, the relative position of the cluster member within the cluster, and the tertiary structure of the cluster all are intricately involved in this regulation (reviewed in Vilimova & Pfeffer, 2023). Our observation within the mammary tumors supports the tertiary structure model of the pri-miR-17-92 carrying a dense supercoiled core with miR-92a being the least accessible by the microprocessor complex (Chaulk et al., 2011). Interesting, this model of the miR-17-92 cluster is different from the tertiary structure proposed by Du and colleagues in which miR-92a is the most accessible member of the cluster for initial processing compared to the remaining cluster members (Du et al., 2015), a model that may apply to the normal mammary epithelial HC11 and HEK293T cells. Thus, the miR-17-92 cluster may adopt a different three-dimensional RNA structure in infected mammary epithelial cells, facilitating the expression of miR-92a, from that of the primary breast tumors in which it may be inhibiting its expression, an aspect that needs to be studied further.

**Figure 8:**
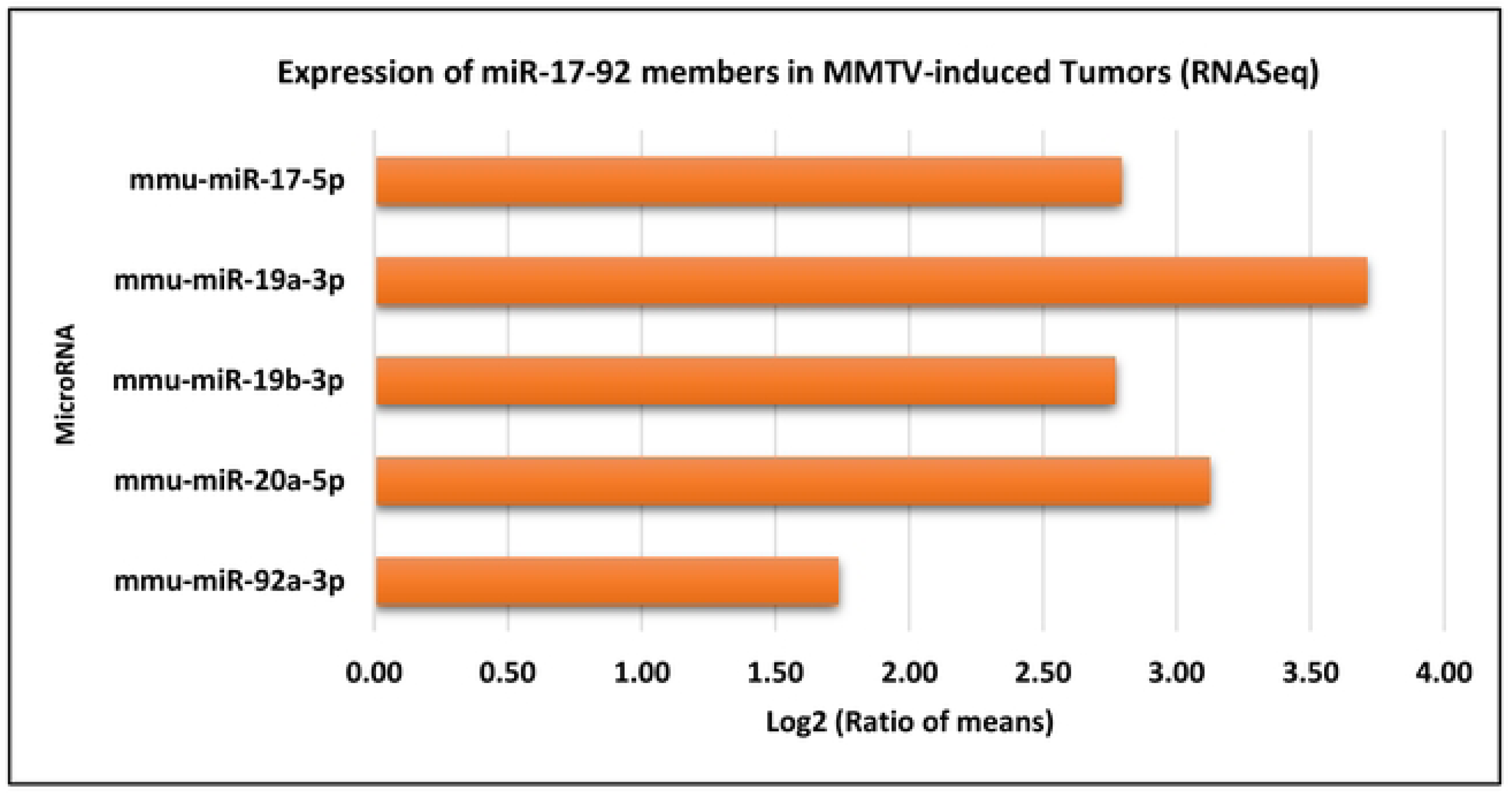
Data representing the mean expression of miR-17-92 cluster members in MMTV-induced tumors (n=2) and infected mammary glands (n=2) obtained by small RNAseq analysis (Kincaid et al., 2018).

Real time qPCR analysis of the MMTV-expressing HEK293T cells revealed the mature miRNA signature of MMTV-infected cells since the expression levels of individual mature cluster members varies, depending upon the infectious agent (Olive et al., 2013, 2015; Fang et al., 2017). A substantial increase in mature miR-92a expression was observed, while levels of other miRs such as miR-17 and miR-19a were not affected significantly (Fig. 1B). Interestingly, over-expression of the miR-17-92 cluster in MMTV-expressing cells led to relatively the same expression profile of the individuals miRs as in MMTV-expressing cells without over-expression of the miR-17-92 cluster, with miR-17 being hardly up-regulated, miR-19a up-regulated by ~ 8-folds, and miR-92a up-regulated the most by ~250-folds (Fig. 2D). The phenotype of miR-92a being the highest expressed member amongst the tested miRs remained in all the three cell lines (Fig. 2D-2F), which is what probably led to the downregulation of MMTV Gag expression in these cell lines (Fig. 2G). This was confirmed by quantitating the suppression of MMTV at the RNA level in all the three miR-over-expressing cell lines in which the level of MMTV mRNA suppression was equivalent to the levels in miR-92a expressing cells alone (Fig. 2I). Thus, the suppression of MMTV expression could be attributed to the high levels of miR-92a in all the three cell lines (Fig. 2H). This data suggests that among the cluster members, miR-92 is the critical component suppressing MMTV expression in cell lines, while its reduced expression in the mammary gland may be responsible for the high levels of virus replication in this tissue, ultimately leading to mammary tumors.

Differential regulation of cluster members by viral infections has been observed before with important biological consequences. For instance, miRNA profiling of blood samples from infants infected by RSV demonstrated reduced levels of miR-92 and increased expression of cluster members miR-20a and paralogue miR-106b-5p, implying their roles in the immune regulation in RSV infections (S. Wang et al., 2017). In our study, the expression of miR-19a was down-regulated in the miR-19/20 OE cell line compared to the empty vector control (Fig. 2E). This was unexpected and could be attributed to known antagonistic interactions between the miR-19 family and miR-92a (Mu et al., 2009; Olive, Sabio, et al., 2013). Olive et al. reported that miR-92 has complex, cell context-dependent roles. In B-cell lymphomas driven by an aberrant expression of c-Myc, miR-92 neutralizes the tumorigenic effect of miR-19 in the presence of p53 by facilitating apoptosis via repression of Fbw7 and thereby stabilizing c-Myc (Olive, Sabio, et al., 2013). The study authors hypothesized that the impairment of the delicate balance between miR-92 and miR-19 levels, with the acquisition of a p53 mutation, fostered the establishment of c-Myc-induced tumors. Since the expression of miR-92a was observed to be significantly up-regulated in HEK293T-MMTV-miR-19/20 cell line (Fig. 2E), it could potentially play a role in modulating the expression of miR-19a. The latter may explain why MMTV Gag expression in the miR-19/20 OE cell line was down-regulated at both the RNA (Fig. 2H) and protein levels (Fig. 2G), even when the miRNAs in question were not over-expressed (Fig. 2D).

With Gag protein levels down-regulated in all three OE cell lines, it was important to determine the mechanism of this downregulation, whether the inhibitory effect was at the level of gRNA expression or its translation into proteins. Test of virions produced by the stable cell lines revealed that the cluster’s inhibitory effect was on viral particle formation since not only were Gag protein levels reduced (Fig. 2J), but also the amount of RNA packaged into the viral particles (Fig. 2I). This could be explained due to the viral gRNA being targeted for degradation in the cells (as seen in Fig. 2I), thereby reducing the availability of Gag proteins and full-length RNA for encapsidation (Fig. 2G). Strikingly, an almost complete absence of MMTV gRNA was observed in the virus particles obtained from miR-92 OE cells (Fig. 2I), highlighting the potency of the inhibitory effect of miR-92a.

By suppressing distinct members of the miR-17-92 cluster independently, miR-17, miR-19a and miR-92a, we established that the suppression of MMTV gene expression could be restored by blocking the repressive miRNAs (Fig. 3). The addition of miRNA inhibitors operate at the mature miRNA level and abolish their ability to bind to the specified target; hence, de-repressing their expression, as detailed by Stenvang et al., 2012. As we used three individual miRNA overexpression cell lines, we aimed at restoring MMTV expression in all three cell lines utilizing anti-miR inhibitors for the corresponding cell line. A combinatorial approach was employed to inhibit three miRNAs in the miR-17-92 OE cell line: miR-17, miR-19a, and miR-92a. Although individual miRNA inhibition was unsuccessful at the miRNA level (Fig. 3A), we noticed a statistically significant increase in MMTV gRNA expression (Fig. 3E) and a modest rescue in Gag protein expression (Fig. 3D). We believe that the non-significant inhibition of target miRNAs in the miR-17-92 OE and miR-19/20 OE cell lines (Fig. 3A & 3B) is probably due to compensatory effects by other cluster members. The observation that miR-92a inhibition in miR-92a OE cells was the highest among other cell lines (Fig. 3C) supports this notion since miR-92a does not share its seed sequence with any other member of the cluster (Grimson et al., 2007; Lewis et al., 2003). Since the repressive miRNAs were inhibited, no significant variation was observed in gRNA levels between the miR-19/20 and miR-92a OE cell lines when compared to the control (Fig. 3F & 3G). However, a statistically significant increase in the gRNA expression was observed in miR-17-92 OE cells (Fig. 3E). We speculate that the miR-92a suppression of MMTV in miR-17-92 OE cells is removed upon the addition of the anti-miR cocktail that contains 15 pmol of anti-miR-92, even though the reduction in miR-92a was statistically non-significant (Fig. 3A). As seen earlier in Fig. 2D, we know that miR-92a is highly expressed in this cell line, thereby the inhibitor, even if active partially, relieves this suppression to a minor extent which was reflected at the gRNA level (Fig. 3E). This was confirmed by a significant rescue of MMTV gene expression upon inhibiting miR-92a at various dosages (Fig. 3H). A considerably better rescue was observed at 15 pmol of inhibitor than at 45 or 60 pmol due to possible saturation of the miRNA binding sites. This notion is in agreement with the work of Mayya and Duchaine, who demonstrated that only a small fraction of miRNAs, determined by a combination of expression threshold, miRISC abundance, and low target site availability, are vulnerable to competitive effects via miRNA-binding sites (Mayya & Duchaine, 2015). However, the role of other cluster members in modulating MMTV replication needs to be explored further since they share seed sequences which may alter their individual effect by collaborative interactions among themselves.

The use of plasmid-based inhibitor vectors aided in the validation of the findings in a cell line system stably expressing anti-miRNAs against miR-17 and miR-92a. Although the PMIS cell lines showed no significant inhibition of target miRNAs, the downstream effect of miRNA suppression was demonstrated at the target level (PTEN), in the case of the representative cell line, PMIS 17 (Fig. 4B). This observation is in line with the published study that developed these plasmids, stating that PMIS molecules form a stable complex with mature miRNAs, thereby preventing their functional role of targeting mRNA down-regulation (Cao et al., 2016). Compared to miR-17, we observed an up-regulation of miR-92a in the PMIS 92 cell line instead of its inhibition (Fig. 4A). We postulate that this observation of no inhibition/up-regulation of target miRNAs is due to significant expression of anti-miRNA molecules in the cell, which is transformed into cDNA and measured in RT-qPCR reactions alongside the miRNAs being quantified (Thomson et al., 2013). Thus, despite the inability to show a physical down-regulation of the targeted miRNAs via PMIS using RT-qPCR assays, we demonstrate a functional inhibition of miR-17 and miR-92a, as observed by a substantial rescue of MMTV Gag at both the RNA and protein levels (Fig. 4D & 4E).

From the inhibition studies presented in Figures 3 and 4, we demonstrate that miR-92a is a powerful anti-viral component that by itself lowers MMTV gene expression to the same level as the entire cluster. Furthermore, we demonstrate that when the miR-92a seed sequence was altered or deleted, MMTV gene expression was restored to levels comparable to the empty vector control (Fig. 5), suggesting a direct interaction of miR-92a with the MMTV genome. Therefore, all prospective miR-17-92 binding sites on the MMTV genome were predicted (Fig. 6). Interestingly, the miRNA binding sites with the highest probability and thermodynamic stability were found in the *gag* region compared to other genes on the viral genome (Fig. 6B). Amongst the predicted thermodynamically stable sites of miR-92, 10 sites were predicted to be concentrated in *gag*, a region that is part of the unspliced mRNA used both for translation of the Gag/Pro/Pol structural/enzymatic proteins as well as encapsidation into the viral particles as the genomic RNA, thus targeting virus replication directly (Fig. 6A). To confirm if these sites were actual targets of the miR-17-92 cluster and miR-92a, luciferase-based assays were used to determine their potential to directly target post-transcriptional silencing of MMTV. We found that the degree of reduction in luciferase activity was nearly the same in all the three OE cell lines (Fig. 7D), most likely due to the high levels of miR-92a expression in these cell lines (Fig. 2D-2F), supporting our assertion that miR-92a directly targets the MMTV genome to induce its suppressive effect. It is critical to note that a specific miRNA-binding site on *gag* can only accommodate a single miRNA at any given time (Arvey et al., 2010), thereby making the target gene susceptible to miRNA-mediated repression regardless of the amounts of exogenous miRNAs. This may explain why we did not observe a dose response when anti-miR-92a oligos were used for inhibition (Fig. 3H) and luciferase activity was down-regulated to about the same extent across the five cell lines assessed (Fig. 7B-7D).

These observations lead to the question of how the miR-17-92 cluster is activated upon MMTV infection? Literature reports transcription factors such as Myc, Stat3, Ccnd1, and members of the E2f family, such as E2f-1 and E2f-3 playing a role in regulating the cluster expression (Brock et al., 2009; Mestdagh et al., 2010; O’Donnell et al., 2005; Sylvestre et al., 2007; Woods et al., 2007). Further experiments are required to investigate the expression of these activators to determine whether any of them are induced upon MMTV expression as a means of activating the miR-17-92 cluster. Our mRNAseq data from MMTV-expressing HC11 cells point to the potential role of c-Myc in this process which had a ~ 3-fold induction in expression in the presence of MMTV, while a ~2-fold down-regulation of Ccnd1 was observed. These observations suggest that c-Myc may be one of the factors involved in the regulation of the miR-17-92 cluster in our system (Supplementary Table S1). Other cellular or viral factors involved in the regulation of the cluster remain to be investigated.

The anti-viral and pro-viral activity of miR-17-92 has been reported sporadically across different viruses and proves to be a promising domain in virology and miRNA-based therapeutics (Rupaimoole & Slack, 2017). For example, its anti-viral potential was demonstrated in HBV infection where c-Myc-induced activation of the miR-17-92 cluster led to inhibition of HBV replication in hepatoma cells (Jung et al., 2013). In the case of enterovirus-71 (EV-71), the virus subverts the anti-viral activity of the cluster through methylation of the cluster promoter, thereby facilitating viral replication and aiding EV-71 infection (Fu et al., 2019). Conversely, the pro-viral activity of the cluster was demonstrated in Kaposi’s sarcoma-associated herpesvirus (KSHV) infection where the cluster was activated by KSHV latency-associated genes, vCyclin and vFLIP. Consequently, this led to downregulation of the immunoregulatory TGF-pathway by targeting SMAD2 suppression, facilitating the promotion and progression of tumors in Kaposi’s sarcoma (Choi et al., 2015). Triboulet et al. reported one of the first studies to show the anti-viral potential of miR-17-92 in retroviruses, revealing that expression of the cluster members was inhibited in HIV-infected Jurkat cells (Triboulet et al., 2007). According to their findings, the cluster indirectly suppressed virus replication by targeting the histone acetylase, PCAF, a key facilitator of transcriptional activation by HIV-1 Tat. Bioinformatic analysis predicted four target sites for miR-17 and miR20a on the 3’ untranslated region (UTR) of the PCAF transcript. This anti-viral role was demonstrated to be effective in combating HIV-1 infection using a multiplex miRNA technology that engineered conserved anti-HIV sequences into the miR-17-92 polycistron, thereby showing its therapeutic potential to combat elusive viral pathogens (Liu et al., 2008).

With the evidence presented in this study, we suggest a negative regulatory interaction between the host miR-17-92 cluster and MMTV whereby MMTV infection and integration into the host genome activates the miR-17-92 cluster. This results in up-regulation of the cluster member miR-92a which directly targets the MMTV genomic RNA, resulting in down-regulation of MMTV Gag/Pro/Pol gene expression as well as nascent virus production (Fig. 9). Hence, our data identifies an anti-viral role of the miR-17-92 cluster in MMTV biology where it negatively regulates the expression of viral genomic RNA, thus inhibiting MMTV replication and *gag* gene expression. In addition to the miR-17-92 cluster down-regulating viral gene expression, cluster member miR-92a was identified as having have an equally strong, negative regulatory effect. This observation points to miR-92a, potentially being the major anti-viral component of the miR-17-92 cluster in regulating MMTV replication. Thus, this study provides the first evidence demonstrating the direct physiological role of miRNAs in modulating MMTV replication. The fact that we observed a downregulation of miR-92a expression in infected mammary glands and mammary tumors *in vivo* further suggests that MMTV is able to disrupt this miR-92a-based negative regulation, most likely via modulating the tertiary structure of the cluster, resulting in enhancing its expression in mammary glands and leading to mammary tumors. This suggests an alternative means of inducing tumorigenesis in addition to the well described process of insertional mutagenesis, an aspect that needs to be explored further.

**Figure 9:**
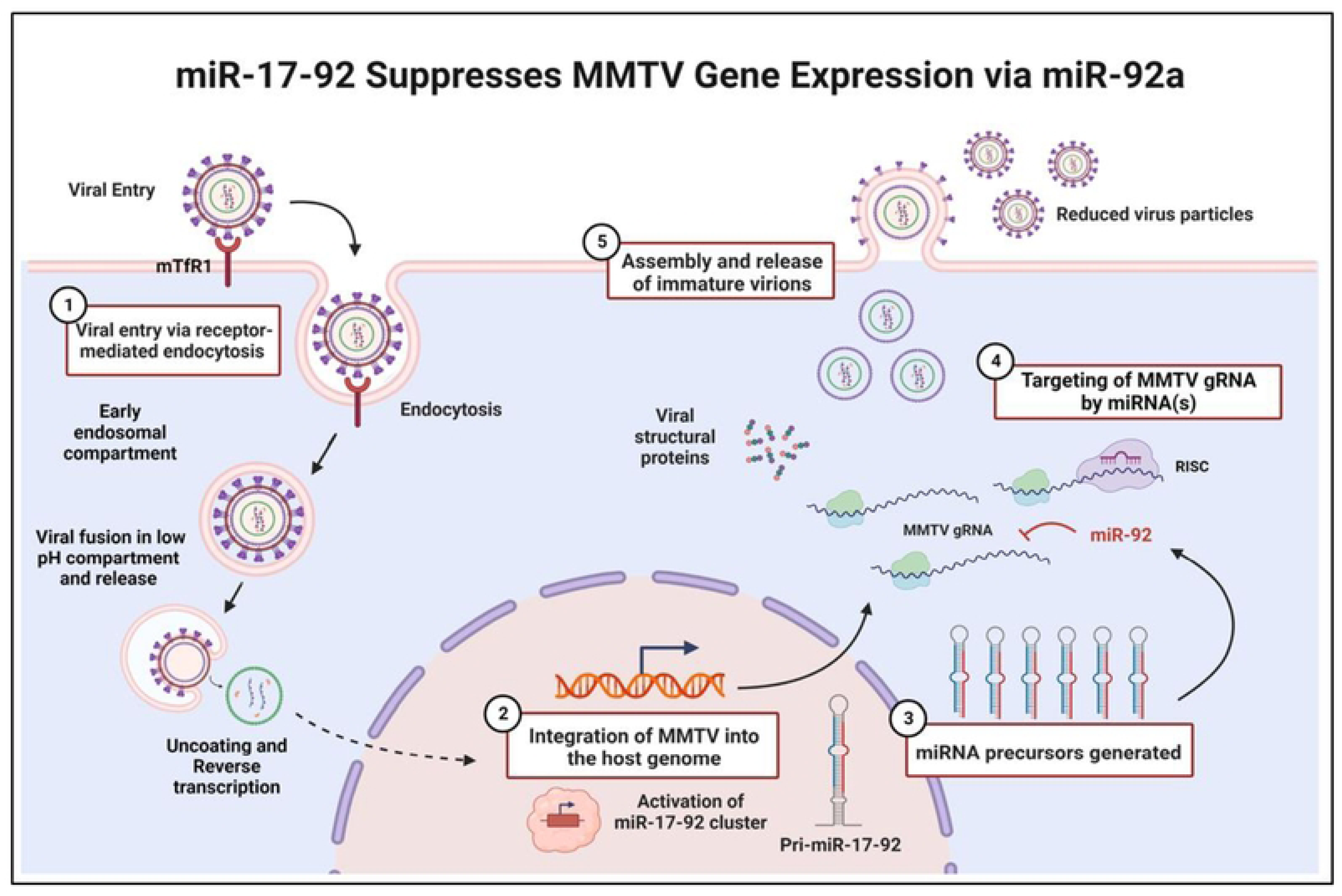
Schematic representation of model representing miR-17-92-mediated regulation of MMTV gene expression. MMTV entry via murine transferrin receptor 1 (mTfR1) is followed by fusion of the viral capsid in a late endosomal compartment under low pH, uncoating/reverse transcription, and integration of the viral genome into the host chromosome. The cellular/viral factor-assisted transcriptional activation of the miR-17-92 cluster leads to up-regulation of the pri-miR-17-92. Once activated, due to the differential processing of the cluster, there is an increase in the expression levels of the mature miR-92a cluster member, which in turn, targets the MMTV unspliced genomic RNA. With more than 10 statistically significant miR-92a binding sites on the viral *gag* region, down-regulation of the genomic RNA takes place, resulting in suppression of MMTV replication. This is because the unspliced genomic RNA serves as the mRNA for the translation of the viral structural and enzymatic proteins (Gag/Pro/Pol) as well as the source of genomic RNA for encapsidation into the newly forming viral particles. Illustration created on *BioRender*.

## Materials and Methods

### Nucleotide Numbering

All nucleotide numbers in this study refer to the nucleotide positions of the exogenous C3H proviral strain of MMTV that is 9,895 bp in length (GenBank accession number AF228552).

### Cell Lines and Culture Conditions

The HEK293T cells were cultured in Dulbecco’s modified Eagle’s medium, DMEM (HyClone, USA), which included 10% fetal bovine serum (HyClone Laboratories Inc., Logan, UT USA). All cell lines had their culture medium supplemented with 1% penicillin and streptomycin (10,000 g/ml; Life Technologies, Carlsbad, CA USA), as well as 0.1% gentamicin (50 mg/ml; Life Technologies, USA). All cell lines were maintained at 37°C and 5% CO_2_ inside a water-jacketed incubator (Forma-series II, ThermoFisher Scientific, USA).

### MMTV Stable Cell Lines

An HEK293T stable cell line that constitutively expressed MMTV was established using HYBMTV, a molecular clone of MMTV that is capable of infection and tumorigenesis (Shackleford & Varmus, 1988). The cells were transfected with linearized HYBMTV plasmid using Lipofectamine 3000 (Invitrogen, ThermoFisher Scientific, USA) according to manufacturer’s instructions. After a period of 48 hours post transfection, the HEK293T-MMTV stable cells were selected for positive clones in hygromycin selection media (200 µg/ml). After four weeks of selection, the isolated colonies were tested for the expression of MMTV Gag by western blotting analysis. The HEK293T (control) cell line was maintained in DMEM and the established HEK293T-MMTV stables were maintained in DMEM-Hygromycin media containing (200 µg/ml). All MMTV expressing cell lines were treated with 10-6 M Dexamethasone (Sigma-Aldrich, Saint Louis, MI USA) 8 hours prior to harvesting to increase MMTV gene expression through hormonal stimulation of its promoter activity, as described by Morley et al., 1987. HC11 cells expressing HYBMTV stably have been described before (Ahmad et al., 2023).

### Plasmids

The miRNA over-expression plasmids and miR-92a mutation/deletion plasmids were obtained from Addgene vector repository, USA (Table 1). The Dual luciferase miRNA target vector (pmirGLO) was purchased from Promega, Madison, WI, USA. The AK14 plasmid used for the construction of the MMTV Gag-containing dual luciferase construct, GagGLO, has been described before (Chameettachal et al., 2018). To test the miR17-92 cluster target sites predicted within Gag, the MMTV Gag expression plasmid, AK14, was digested with *Xho*I to obtain the complete MMTV Gag region (nt 1485-nt 3258). The *gag* gene was then cloned into the *Xho*I site downstream of the *firefly luciferase* gene of pmirGLO to create the Gag target vector, GagGLO. All constructs used in the study were verified by sequencing and restriction digestion and are listed in Table 1.

**Table 1:** List of plasmids used in this study.

| Plasmid | Function | Source | Reference |
| --- | --- | --- | --- |
| MSCV | Empty vector control with GFP and Puromycin selectable markers | Addgene (#24828) | Olive et al., 2009 |
| MSCV-miR-17-92 | MSCV vector expressing wild type miR-17-92 cluster | Addgene (#64100) | Olive, Sabio, et al., 2013 |
| MSCV-miR-19a-20a-19b | MSCV vector expressing miR-19a, miR-20a and miR-19b of the cluster | Addgene (#24827) | Olive et al., 2009 |
| MSCV-miR-92a | MSCV vector expressing miR-92a of the cluster | Addgene (#64092) | Olive et al., 2009 |
| PMIS NC | Control vector with scramble oligo | NaturemiRI | H. Cao et al., 2016 |
| PMIS17 | PMIS vector expressing anti-miR-17 | NaturemiRI | H. Cao et al., 2016 |
| PMIS92a | PMIS vector expressing anti-miR-92a | NaturemiRI | H. Cao et al., 2016 |
| MLS-miR-17-92a | MLS vector expressing wild type miR-17-92 cluster | Addgene (#64090) | Olive et al., 2009 |
| MLS-miR-17-19b | MLS vector expressing truncated miR-17-19b cluster w/o miR-92a | Addgene (t#64089) | Olive et al., 2009 |
| miR-17-92aMUT | MLS vector expressing miR-17-92 cluster w/ mutated miR-92a | Addgene (#64094) | Olive, Sabio, et al., 2013 |
| AK14 | Gag expression vector | Tahir A. Rizvi | Chameettachal et al., 2018 |

### miRNA Over-expression and Inhibition Experiments

For miRNA over-expression or plasmid-based inhibition analysis, the HEK293T-MMTV cells were transfected with the appropriate plasmids (listed in Table 1) using Lipofectamine 3000 (Invitrogen, ThermoFisher Scientific, USA), as described by the manufacturer. Briefly, 5×10^5^ cells were plated per well of a 12-well plate the day before transfection. The plasmid DNA (3μg/well) was added to the DNA/lipid cocktail, followed by incubation for 15 minutes at room temperature. The mixture was added to the cells dropwise while swirling. The transfected cells were returned to the incubator and inspected for GFP positive cells at the 48-hour time point. If positive, they were selected in 3 μg/μl puromycin or another appropriate antibiotic selection media for 5 or more days for stable selection.

To prepare viral particles, cell supernatant was pre-cleared by low-speed centrifugation followed by filtering through a 0.2µ filter. Virus particles from the clarified cell supernatants were purified by ultra-centrifugation at 26,000 rpm using the SW50.1 rotor for 90 minutes through a 2 ml, 20% sucrose cushion. The viral pellets obtained were then scrapped into 300 μl of 1× TNE buffer (1 mM EDTA, 10 mM Tris-HCl of pH 7.5, 100 mM NaCl) and divided for RNA and protein analyses.

### miRNA Inhibition Experiments using Anti-miR Oligos

Functional inhibition of miRNAs in the cluster was achieved using the potent mirVana oligo-based inhibitors for miR-17, miR-19a and miR-92a (ThermoFisher Scientific, USA): miR-17 (MIMAT0000070), miR-19a (MIMAT0000073), and miR-92a (MIMAT0000092) along with negative control (NEG#1 Cat.No:4464076). The oligo inhibitors were resuspended in nuclease free water to make 100 μM inhibitor stock solution, according to the manufacturer’s protocol. Inhibitors (alongside a negative scramble control) were used at a final concentration of 15 pmol/well and were transfected into the specific over-expression cell line using Lipofectamine 3000. A combination of miR-17, miR-19, and miR-92a inhibitors (15 pmol/inhibitor each) was used to treat the complete miR-17-92 cluster over-expression cell line with the intention of inhibiting the three miRNAs in the cell line simultaneously. The treated cells were harvested 72 hours post transfection and processed for RNA and protein for further analysis.

### RNA Sequencing (mRNAseq and miRNAseq)

Total cellular RNA was extracted from normal mouse mammary epithelial HC11 cells or HC11 cells expressing MMTV, HC11-MMTV, using the TRIzol Reagent (Invitrogen, ThermoFisher Scientific, Waltham, MA USA). Whole cell RNA was sequenced for both mRNA and miRNA expression from two independent biological replicates of the HC11 and HC11-MMTV cell lines using the TruSeq library-NovaSeq6000 platform (for mRNAseq) or TruSeq SBS KIT-HS V3-HiSeq 2000 system (for small RNAseq) commercially by the Beijing Genomics Institute (BGI, Hong Kong). Data analysis was carried out using BGI’s online tool, Dr. Tom. The validated mRNAseq data has already been published (Ahmad et al., 2023).

### RNA Extraction and cDNA Synthesis

RNA was extracted from cells and viral particles using the Trizol Reagent (ThermoFisher Scientific, USA) and quantified using a Nanodrop spectrophotometer. To ensure DNA-free preparations, 6 μg of the extracted RNA was DNase-treated for 30 minutes at 37°C with 2 units (U) of Turbo DNase I (Invitrogen, ThermoFisher Scientific, USA) along with 40 units of Recombinant RNasin (Promega, USA). After confirmation of DNA free preparations using PCR, the RNA was further reverse transcribed to cDNA using MMLV-RT (Promega, USA) and used for downstream mRNA qPCR assays. For miRNA quantification, 10 ng of isolated total RNA was converted into cDNA using the TaqMan™ miRNA Reverse Transcription Kit (Applied Biosystems, Foster City, CA, USA) which was subsequently used for stem-loop-based real time PCR using TaqMan® miRNA Individual Assays and the TaqMan® Universal PCR Master Mix with AmpErase® UNG (Applied Biosystems, USA) (Chen et al., 2005).

### Quantitation of MMTV by Real-time PCR

Real-time quantitation of MMTV full length genomic and all mRNAs was performed using customized TaqMan assays (FAM-labelled) described in Mustafa et al., 2012; Aktar et al., 2014. Briefly, all MMTV mRNAs were quantified using an assay specific for the 5’ U5 region within the HYB-MTV genome (nt 1192-1259) and genomic RNA was quantified using an assay that targeted the nucleotide region 1729-1791 of MMTV *gag*. The experiments were normalized using human β-Actin as the endogenous control (FAM/MGB probe, cat. no. 401846; Applied Biosystems, USA).

### Quantitation of Pri-miRNAs by SYBR Green qPCR

The pri-form of miR-17-92 cluster was quantified using SYBR Green qPCR assay utilizing 5X HOT FIREPol® EvaGreen® qPCR Mix (Solis BioDyne, Estonia) with two individual primer sets described earlier (Chakraborty et al., 2012; Z. Wang et al., 2010) that were renamed OFM 450/451, OFM 446/447 in this study. The amplification reaction contained template nucleic acid (1 μl of cDNA), 4 μl 5X HOT FIREPol® EvaGreen® qPCR mix, 0.8 μl per Primer (400 nM), and nuclease-free water in a total reaction volume of 20 μl. The qPCRs were performed in triplicates under the following cycling conditions: 50°C for 2 mins, initial denaturation at 95°C for 12 mins, 40 cycles of 95°C for 15 secs, annealing at 60°C for 30 secs, extension 72°C for 30 secs, and a final extension at 72°C for 10 min. The primers used for the assays are listed in Table 2.

**Table 2:**
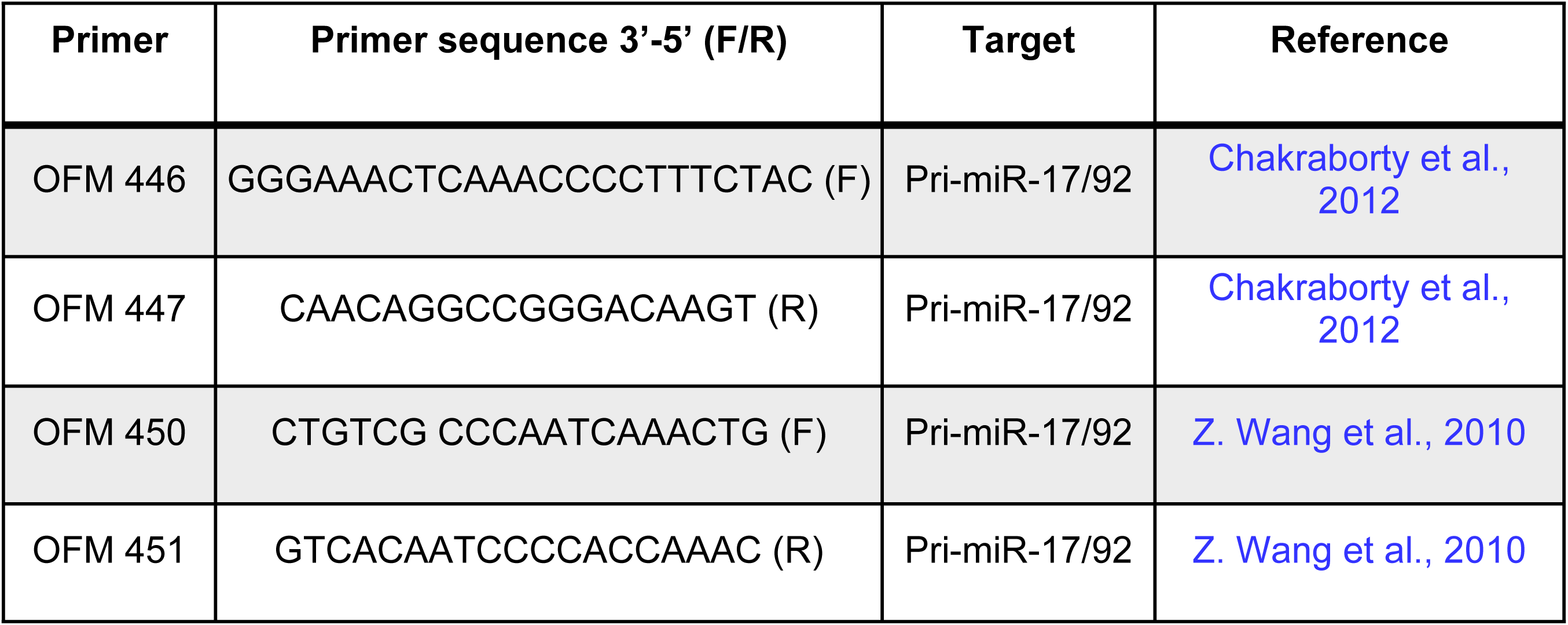
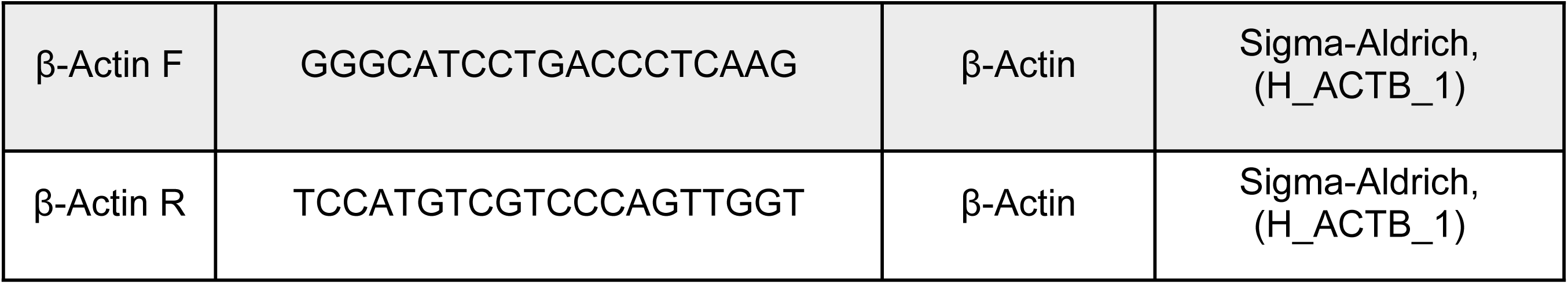
Primers used to quantify pri-miR-17-92 and the endogenous control β-Actin.

### Quantitation of Mature miRNAs by TaqMan Assays

The mature miRNA species were quantified using the previously-validated TaqMan miRNA assays for miR-17 (Assay ID: 002308), miR-19a (Assay ID: 000395) and miR-92a (Assay ID:000430) (Applied Biosystems, USA), as per manufacturer’s protocol. The experiment was normalized using U6 snRNA (Assay ID: 001973) as the endogenous small RNA control. The assays were carried out in a volume of 20 μl in triplicates using QuantStudio™7 Flex Real-Time PCR System following the standard conditions: 2 minutes at 50°C, 10 minutes at 95°C for denaturation, followed by 40 cycles of denaturation at 94°C for 15 seconds and annealing/extension for 1 minute at 60°C. The data was analyzed using the 2^−ΔΔCt^ method (Livak & Schmittgen, 2001).

### Western Blotting

Whole cell extracts and virus particles were analyzed by western blotting to quantify MMTV protein expression. Harvested cells were lysed in RIPA buffer [1 mM EDTA, 10 mM Tris–HCl (pH 8.0), 0.1% sodium deoxycholate, 1% Triton X-100,140 mM NaCl and 0.1% SDS] supplemented with 50 μl β-mercaptoethanol and 1 mM PMSF (Sigma-Aldrich, USA) per ml of RIPA buffer. For the analysis of extracellular viral particles, a volume of 50 μl of 1X TNE solution was used to resuspended the virions pellet by ultracentrifugation followed by addition of the 6x SDS loading buffer to 1x and SDS-PAGE separation. Extracted cellular proteins were quantified using the Bradford Reagent (Bio-Rad Laboratories, Hercules, CA USA) as per manufacturer’s directions. Subsequently, lysates were separated on 4-12% SDS polyacrylamide gels and transferred overnight at 4°C, 30 V or at 90 minutes, 90V onto nitrocellulose membranes (GE Healthcare, Chicago, IL USA). The blocked membranes were probed with either rabbit polyclonal MMTV anti-Gag antibody (Rockland Immunochemicals, Inc, Limerick, PA USA) at a dilution of 1:1000 in 2% non-fat dry milk overnight at 4°C, or anti-GAPDH followed by a 1-hour incubation at room temperature with the appropriate secondary antibody conjugated with horseradish peroxidase. The chemiluminescent signal was detected using the ECL Plus Western blotting substrate or SuperSignal™ West Femto Maximum Sensitivity Substrate (ThermoFischer Scientific, USA), as directed by the manufacturer, and captured using the TyphoonFLA9500 (GE Healthcare, USA) or an X-ray film. GelQuantNET software was employed for densitometric analysis.

### Dual Luciferase Assays

For miRNA target analysis, HEK293T, HEK293T-MMTV and HEK293T cell lines over-expressing miR-17-92, miR-19a/20a, and miR-92a (1×10^5^ cells in 24-well plates) were transfected with the Dual Luciferase constructs-pmirGLO or the MMTV-GAG containing, GagGLO using Lipofectamine 3000, as described by the manufacturer. The cells were harvested 48 hours post transfection, and whole cell lysates prepared and assayed for luciferase activity using the Dual Luciferase Assay System as per manufacturer’s directions (Promega, USA). The luminescence signal was detected using the luminometer (Promega). The Firefly luciferase activity was normalized to Renilla luciferase expression and expressed relative to the experimental control, pmirGLO. All assays were conducted in triplicates and independently repeated two times.

### miRNA-mRNA Target Prediction

All potential miR-17-92 binding sites on the MMTV genome were predicted using Sfold StarMirDB web server for miRNAs: (https://sfold.wadsworth.org/starmirDB.php) (Kanoria et al., 2016; Rennie et al., 2019). The following parameters were followed through the bioinformatic analysis: Logistic probability > 0.75, ΔGHybrid ≤ −20kcal/mol, Preferential Seed type −8mer/7mer (Rennie et al., 2016).

### Statistical Analysis

Statistical analysis was performed using GraphPad Prism Version 7.0. Statistical significance was determined using the paired student’s *t* test, as the mean ± SD. All recorded data in this study were acquired from at least 2 independent experiments. A *p* value of 0.05 was considered to be least significant where *p is ≤ 0.05; **p ≤ 0.01, ***p ≤0.001, ****p≤ 0.0001. ns (not significant), p > 0.05.

To determine significance between groups in small RNAseq data, Q value was estimated which is basically the *p* value adjusted for the false discovery rate (FDR) using the multiple hypothesis test. The closer the Q value is to zero, the more significant is the data (log2 ratio in our case) between two groups.

## Acknowledgments

We would like to thank Prof. Jeffery M. Rosen, Baylor College of Medicine, Houston, TX USA for the gift of HC11 cells, and the Lin He Lab (Lin He, Professor of Cell Biology, Development & Physiology University of California, Berkeley CA, USA) for sharing the miRNA overexpression plasmids: MSCV, MSCV-mir-17-92, MSCV-19a-20-19b, MSCV-mir-92, MLS-mir-17-92, MLS-mir-17-19b, MLS-mir-17-92-Mut92 (Addgene plasmid #: 24828, 64100, 24827, 64092, 64090, 64089, & 64094). We are also grateful to the Amendt Lab (Prof. Brad A. Amendt, Department of Anatomy & Cell Biology, Carver College of Medicine, University of Iowa, Iowa City, IA USA) for providing the PMIS miRNA inhibitor vectors.

## Author Contributions

**Conceptualization:** FM, JB, BG**; Data curation:** JB, BG, WA**; Formal analysis:** JB, BG, WA; **Funding acquisition:** FM; **Investigation:** JB, WA, BG, NGP, TAK, SQ; **Methodology:** JB, BG, WA, NGP, TAK, SQ; **Project administration:** FM; **Resources:** FM, TAR; **Software:** JB, BG, WA; **Supervision:** FM; **Validation:** JB, WA; **Visualization:** JB, WA, FM; **Writing-original draft:** JB; **Writing-review & editing:** JB, TAR, FM,.

## References

Ahmad, W., Panicker, N. G., Akhlaq, S., Gull, B., Baby, J., Khader, T. A., Rizvi, T. A., & Mustafa, F. (2023). Global down-regulation of gene expression induced by mouse mammary tumor virus (MMTV) in normal mammary epithelial cells. Viruses 15(5), 1110. 10.3390/v15051110.

Aktar, S. J., Vivet-Boudou, V., Ali, L. M., Jabeen, A., Kalloush, R. M., Richer, D., Mustafa, F., Marquet, R., & Rizvi, T. A. (2014). Structural basis of genomic RNA (gRNA) dimerization and packaging determinants of mouse mammary tumor virus (MMTV). Retrovirology, 11(1), 96. 10.1186/s12977-014-0096-6

Amarante, M. K., de Sousa Pereira, N., Vitiello, G. A. F., & Watanabe, M. A. E. (2019). Involvement of a mouse mammary tumor virus (MMTV) homologue in human breast cancer: Evidence for, against and possible causes of controversies. Microbial Pathogenesis, 130, 283–294. 10.1016/j.micpath.2019.03.021

Arvey, A., Larsson, E., Sander, C., Leslie, C. S., & Marks, D. S. (2010). Target mRNA abundance dilutes microRNA and siRNA activity. Molecular Systems Biology, 6, 363. 10.1038/msb.2010.24

Baby, J. (2022). The Host miR-17-92 Cluster Negatively Regulates MMTV Replication by Targeting its Genomic RNA via miR-92a. Doctoral Thesis. United Arab Emirates (UAE) University.

Bennasser, Y., Le, S.-Y., Yeung, M. L., & Jeang, K.-T. (2004). HIV-1 encoded candidate micro-RNAs and their cellular targets. Retrovirology, 1(1), 43. 10.1186/1742-4690-1-43

Bevilacqua, G. (2022). The Viral Origin of Human Breast Cancer: From the Mouse Mammary Tumor Virus (MMTV) to the Human Betaretrovirus (HBRV). Viruses, 14(8), Article 8. 10.3390/v14081704

Brock, M., Trenkmann, M., Gay, R. E., Michel, B. A., Gay, S., Fischler, M., Ulrich, S., Speich, R., & Huber, L. C. (2009). Interleukin-6 Modulates the Expression of the Bone Morphogenic Protein Receptor Type II Through a Novel STAT3–microRNA Cluster 17/92 Pathway. Circulation Research, 104(10), 1184–1191. 10.1161/CIRCRESAHA.109.197491

Cao, H., Yu, W., Li, X., Wang, J., Gao, S., Holton, N. E., Eliason, S., Sharp, T., & Amendt, B. A. (2016). A new plasmid-based microRNA inhibitor system that inhibits microRNA families in transgenic mice and cells: A potential new therapeutic reagent. Gene Therapy, 23(6), 527–542. 10.1038/gt.2016.22

Chakraborty, S., Mehtab, S., Patwardhan, A., & Krishnan, Y. (2012). Pri-miR-17-92a transcript folds into a tertiary structure and autoregulates its processing. RNA, 18(5), 1014–1028. 10.1261/rna.031039.111

Chameettachal, A., Pillai, V. N., Ali, L. M., Pitchai, F. N. N., Ardah, M. T., Mustafa, F., Marquet, R., & Rizvi, T. A. (2018). Biochemical and Functional Characterization of Mouse Mammary Tumor Virus Full-Length Pr77Gag Expressed in Prokaryotic and Eukaryotic Cells. Viruses, 10(6), 334. 10.3390/v10060334

Chaulk, S. G., Thede, G. L., Kent, O. A., Xu, Z., Gesner, E. M., Veldhoen, R. A., Khanna, S. K., Goping, I. S., MacMillan, A. M., Mendell, J. T., Young, H. S., Fahlman, R. P., & Glover, J. N. M. (2011). Role of pri-miRNA tertiary structure in miR-17~92 miRNA biogenesis. RNA Biology, 8(6), 1105–1114. 10.4161/rna.8.6.17410

Chen, C., Ridzon, D. A., Broomer, A. J., Zhou, Z., Lee, D. H., Nguyen, J. T., Barbisin, M., Xu, N. L., Mahuvakar, V. R., Andersen, M. R., Lao, K. Q., Livak, K. J., & Guegler, K. J. (2005). Real-time quantification of microRNAs by stem-loop RT-PCR. Nucleic Acids Research, 33(20), e179. 10.1093/nar/gni178

Choi, H. S., Jain, V., Krueger, B., Marshall, V., Kim, C. H., Shisler, J. L., Whitby, D., & Renne, R. (2015). Kaposi’s Sarcoma-Associated Herpesvirus (KSHV) Induces the Oncogenic miR-17-92 Cluster and Down-Regulates TGF-β Signaling. PLOS Pathogens, 11(11), e1005255. 10.1371/journal.ppat.1005255

Concepcion, C. P., Bonetti, C., & Ventura, A. (2012). The miR-17-92 family of microRNA clusters in development and disease. Cancer Journal (Sudbury, Mass.), 18(3), 262–267. 10.1097/PPO.0b013e318258b60a

Cullen, B. R. (2006). Viruses and microRNAs. Nature Genetics, 38 Suppl, S25–30. 10.1038/ng1793

Cullen, B. R. (2012). MicroRNA expression by an oncogenic retrovirus. Proceedings of the National Academy of Sciences, 109(8), 2695–2696. 10.1073/pnas.1200328109

Czech, B., Malone, C. D., Zhou, R., Stark, A., Schlingeheyde, C., Dus, M., Perrimon, N., Kellis, M., Wohlschlegel, J. A., Sachidanandam, R., Hannon, G. J., & Brennecke, J. (2008). An endogenous small interfering RNA pathway in Drosophila. Nature, 453(7196), 798–802. 10.1038/nature07007

de Pontual, L., Yao, E., Callier, P., Faivre, L., Drouin, V., Cariou, S., Van Haeringen, A., Geneviève, D., Goldenberg, A., Oufadem, M., Manouvrier, S., Munnich, A., Vidigal, J. A., Vekemans, M., Lyonnet, S., Henrion-Caude, A., Ventura, A., & Amiel, J. (2011). Germline deletion of the miR-17~92 cluster causes skeletal and growth defects in humans. Nature Genetics, 43(10), Article 10. 10.1038/ng.915

Ding, S.-W., & Voinnet, O. (2007). Antiviral Immunity Directed by Small RNAs. Cell, 130(3), 413–426. 10.1016/j.cell.2007.07.039

Du, P., Wang, L., Sliz, P., & Gregory, R. I. (2015). A Biogenesis Step Upstream of Microprocessor Controls miR-17~92 Expression. Cell, 162(4), 885–899. 10.1016/j.cell.2015.07.008

Dudley, J. P., Golovkina, T. V., & Ross, S. R. (2016). Lessons Learned from Mouse Mammary Tumor Virus in Animal Models. ILAR Journal, 57(1), 12–23. 10.1093/ilar/ilv044

Fang, L.-L., Wang, X.-H., Sun, B.-F., Zhang, X.-D., Zhu, X.-H., Yu, Z.-J., & Luo, H. (2017). Expression, regulation and mechanism of action of the miR-17-92 cluster in tumor cells (Review). International Journal of Molecular Medicine, 40(6), 1624–1630. 10.3892/ijmm.2017.3164

Flynt, A. S., & Lai, E. C. (2008). Biological principles of microRNA-mediated regulation: Shared themes amid diversity. Nature Reviews Genetics, 9(11), 831–842. 10.1038/nrg2455

Frasca, F., Scordio, M., & Scagnolari, C. (2022). Chapter 15—MicroRNAs and the immune system. In J. Xiao (Ed.), MicroRNA (pp. 279–305). Academic Press. 10.1016/B978-0-323-89774-7.00007-8

Fu, Y., Zhang, L., Zhang, R., Xu, S., Wang, H., Jin, Y., & Wu, Z. (2019). Enterovirus 71 Suppresses miR-17-92 Cluster Through Up-Regulating Methylation of the miRNA Promoter. Frontiers in Microbiology, 10. https://www.frontiersin.org/articles/10.3389/fmicb.2019.00625

Gillet, N., Florins, A., Boxus, M., Burteau, C., Nigro, A., Vandermeers, F., Balon, H., Bouzar, A.-B., Defoiche, J., Burny, A., Reichert, M., Kettmann, R., & Willems, L. (2007). Mechanisms of leukemogenesis induced by bovine leukemia virus: Prospects for novel anti-retroviral therapies in human. Retrovirology, 4(1), 18. 10.1186/1742-4690-4-18

Girardi, E., López, P., & Pfeffer, S. (2018). On the Importance of Host MicroRNAs During Viral Infection. Frontiers in Genetics, 9. https://www.frontiersin.org/articles/10.3389/fgene.2018.00439

Glasgow, L. A. (1970). Cellular Immunity in Host Resistance to Viral Infections. Archives of Internal Medicine, 126(1), 125–134. 10.1001/archinte.1970.00310070127011

Grassmann, R., & Jeang, K. (2008). The roles of microRNAs in mammalian virus infection. Biochimica et Biophysica Acta (BBA) - Gene Regulatory Mechanisms, 1779(11), 706–711. 10.1016/j.bbagrm.2008.05.005

Grimson, A., Farh, K. K.-H., Johnston, W. K., Garrett-Engele, P., Lim, L. P., & Bartel, D. P. (2007). MicroRNA Targeting Specificity in Mammals: Determinants beyond Seed Pairing. Molecular Cell, 27(1), 91–105. 10.1016/j.molcel.2007.06.017

Gull, B, Ahmad, W., Baby, J., Panicker, N. G., Khader, T. A., Akhlaq, S., Rizvi, T. A., Mustafa, F. Characterization of miRNAs that Target the Mouse Mammary Tumor Virus (MMTV) Genome. Submitted.

Ha, M., & Kim, V. N. (2014). Regulation of microRNA biogenesis. Nature Reviews Molecular Cell Biology, 15(8), Article 8. 10.1038/nrm3838

Hamilton, A. J., & Baulcombe, D. C. (1999). A Species of Small Antisense RNA in Posttranscriptional Gene Silencing in Plants. Science, 286(5441), 950–952. 10.1126/science.286.5441.950

He, L., Thomson, J. M., Hemann, M. T., Hernando-Monge, E., Mu, D., Goodson, S., Powers, S., Cordon-Cardo, C., Lowe, S. W., Hannon, G. J., & Hammond, S. M. (2005). A microRNA polycistron as a potential human oncogene. Nature, 435(7043), 828–833. 10.1038/nature03552

Holt, M. P., Shevach, E. M., & Punkosdy, G. A. (2013). Endogenous Mouse Mammary Tumor Viruses (Mtv): New Roles for an Old Virus in Cancer, Infection, and Immunity. Frontiers in Oncology, 3, 287. 10.3389/fonc.2013.00287

Hook, L. M., Agafonova, Y., Ross, S. R., Turner, S. J., & Golovkina, T. V. (2000). Genetics of Mouse Mammary Tumor Virus-Induced Mammary Tumors: Linkage of Tumor Induction to the gagGene. Journal of Virology, 74(19), 8876–8883. 10.1128/JVI.74.19.8876-8883.2000

Houzet, L., & Jeang, K.-T. (2011). MicroRNAs and human retroviruses. Biochimica et Biophysica Acta (BBA) - Gene Regulatory Mechanisms, 1809(11), 686–693. 10.1016/j.bbagrm.2011.05.009

Jung, Y. J., Kim, J.-W., Park, S. J., Min, B. Y., Jang, E. S., Kim, N. Y., Jeong, S.-H., Shin, C. M., Lee, S. H., Park, Y. S., Hwang, J.-H., Kim, N., & Lee, D. H. (2013). C-Myc-mediated overexpression of miR-17-92 suppresses replication of hepatitis B virus in human hepatoma cells. Journal of Medical Virology, 85(6), 969–978. 10.1002/jmv.23534

Kanoria, S., Rennie, W., Liu, C., Carmack, C. S., Lu, J., & Ding, Y. (2016). STarMir Tools for Prediction of microRNA binding sites. Methods in Molecular Biology (Clifton, N.J.), 1490, 73–82. 10.1007/978-1-4939-6433-8_6

Katz, E., Lareef, M. H., Rassa, J. C., Grande, S. M., King, L. B., Russo, J., Ross, S. R., & Monroe, J. G. (2005). MMTV Env encodes an ITAM responsible for transformation of mammary epithelial cells in three-dimensional culture. Journal of Experimental Medicine, 201(3), 431–439. 10.1084/jem.20041471

Kincaid, R. P., Burke, J. M., & Sullivan, C. S. (2012). RNA virus microRNA that mimics a B-cell oncomiR. Proceedings of the National Academy of Sciences, 109(8), 3077–3082. 10.1073/pnas.1116107109

Kincaid, R. P., Chen, Y., Cox, J. E., Rethwilm, A., & Sullivan, C. S. (2014). Noncanonical MicroRNA (miRNA) Biogenesis Gives Rise to Retroviral Mimics of Lymphoproliferative and Immunosuppressive Host miRNAs. MBio, 5(2), e00074–14. 10.1128/mBio.00074-14

Kincaid, R. P., Panicker, N. G., Lozano, M. M., Sullivan, C. S., Dudley, J. P., & Mustafa, F. (2018). MMTV does not encode viral microRNAs but alters the levels of cancer-associated host microRNAs. Virology, 513, 180–187. 10.1016/j.virol.2017.09.030

Kincaid, R. P., & Sullivan, C. S. (2012). Virus-Encoded microRNAs: An Overview and a Look to the Future. PLOS Pathogens, 8(12), e1003018. 10.1371/journal.ppat.1003018

Klase, Z. A., Sampey, G. C., & Kashanchi, F. (2013). Retrovirus infected cells contain viral microRNAs. Retrovirology, 10(1), 15. 10.1186/1742-4690-10-15

Koralov, S. B., Muljo, S. A., Galler, G. R., Krek, A., Chakraborty, T., Kanellopoulou, C., Jensen, K., Cobb, B. S., Merkenschlager, M., Rajewsky, N., & Rajewsky, K. (2008). Dicer Ablation Affects Antibody Diversity and Cell Survival in the B Lymphocyte Lineage. Cell, 132(5), 860–874. 10.1016/j.cell.2008.02.020

Koyama, S., Ishii, K. J., Coban, C., & Akira, S. (2008). Innate immune response to viral infection. Cytokine, 43(3), 336–341. 10.1016/j.cyto.2008.07.009

Le Grice, S. F. J. (2012). Human Immunodeficiency Virus Reverse Transcriptase: 25 Years of Research, Drug Discovery, and Promise. The Journal of Biological Chemistry, 287(49), 40850–40857. 10.1074/jbc.R112.389056

Lewis, B. P., Burge, C. B., & Bartel, D. P. (2005). Conserved Seed Pairing, Often Flanked by Adenosines, Indicates that Thousands of Human Genes are MicroRNA Targets. Cell, 120(1), 15–20. 10.1016/j.cell.2004.12.035

Lewis, B. P., Shih, I.-hung, Jones-Rhoades, M. W., Bartel, D. P., & Burge, C. B. (2003). Prediction of mammalian microRNA targets. Cell, 115(7), 787–798. 10.1016/s0092-8674(03)01018-3

Lin, J., & Cullen, B. R. (2007). Analysis of the Interaction of Primate Retroviruses with the Human RNA Interference Machinery. Journal of Virology, 81(22), 12218–12226. 10.1128/JVI.01390-07

Lindsay, M. A. (2008). MicroRNAs and the immune response. Trends in Immunology, 29(7), 343–351. 10.1016/j.it.2008.04.004

Liu, Y. P., Haasnoot, J., ter Brake, O., Berkhout, B., & Konstantinova, P. (2008). Inhibition of HIV-1 by multiple siRNAs expressed from a single microRNA polycistron. Nucleic Acids Research, 36(9), 2811–2824. 10.1093/nar/gkn109

Livak, K. J., & Schmittgen, T. D. (2001). Analysis of Relative Gene Expression Data Using Real-Time Quantitative PCR and the 2−ΔΔCT Method. Methods, 25(4), 402–408. 10.1006/meth.2001.1262

Mallick, B., Ghosh, Z., & Chakrabarti, J. (2009). MicroRNome analysis unravels the molecular basis of SARS infection in bronchoalveolar stem cells. PloS One, 4(11), e7837. 10.1371/journal.pone.0007837

Mayya, V. K., & Duchaine, T. F. (2015). On the availability of microRNA-induced silencing complexes, saturation of microRNA-binding sites and stoichiometry. Nucleic Acids Research, 43(15), 7556–7565. 10.1093/nar/gkv720

Medley, J. C., Panzade, G., & Zinovyeva, A. Y. (2021). microRNA strand selection: Unwinding the rules. Wiley Interdisciplinary Reviews. RNA, 12(3), e1627. 10.1002/wrna.1627

Mestdagh, P., Boström, A.-K., Impens, F., Fredlund, E., Van Peer, G., De Antonellis, P., von Stedingk, K., Ghesquière, B., Schulte, S., Dews, M., Thomas-Tikhonenko, A., Schulte, J. H., Zollo, M., Schramm, A., Gevaert, K., Axelson, H., Speleman, F., & Vandesompele, J. (2010). The miR-17-92 microRNA cluster regulates multiple components of the TGF-β pathway in neuroblastoma. Molecular Cell, 40(5), 762–773. 10.1016/j.molcel.2010.11.038

Mogilyansky, E., & Rigoutsos, I. (2013). The miR-17/92 cluster: A comprehensive update on its genomics, genetics, functions and increasingly important and numerous roles in health and disease. Cell Death & Differentiation, 20(12), Article 12. 10.1038/cdd.2013.125

Moi, L., Braaten, T., Al-Shibli, K., Lund, E., & Busund, L.-T. R. (2019). Differential expression of the miR-17-92 cluster and miR-17 family in breast cancer according to tumor type; results from the Norwegian Women and Cancer (NOWAC) study. Journal of Translational Medicine, 17(1), 334. 10.1186/s12967-019-2086-x

Morley, K. L., Toohey, M. G., & Peterson, D. O. (1987). Transcriptional repression of a hormone-responsive promoter. Nucleic Acids Research, 15(17), 6973–6989. 10.1093/nar/15.17.6973

Mu, P., Han, Y.-C., Betel, D., Yao, E., Squatrito, M., Ogrodowski, P., de Stanchina, E., D’Andrea, A., Sander, C., & Ventura, A. (2009). Genetic dissection of the miR-17~92 cluster of microRNAs in Myc-induced B-cell lymphomas. Genes & Development, 23(24), 2806–2811. 10.1101/gad.1872909

Mukherjee, A., Di Bisceglie, A. M., & Ray, R. B. (2015). Hepatitis C Virus-Mediated Enhancement of MicroRNA miR-373 Impairs the JAK/STAT Signaling Pathway. Journal of Virology, 89(6), 3356–3365. 10.1128/JVI.03085-14

Mustafa, F., Al Amri, D., Al Ali, F., Al Sari, N., Al Suwaidi, S., Jayanth, P., Philips, P. S., & Rizvi, T. A. (2012). Sequences within both the 5’ UTR and Gag are required for optimal in vivo packaging and propagation of mouse mammary tumor virus (MMTV) genomic RNA. PloS One, 7(10), e47088. 10.1371/journal.pone.0047088

Nahand, J. S., Mahjoubin-Tehran, M., Moghoofei, M., Pourhanifeh, M. H., Mirzaei, H. R., Asemi, Z., Khatami, A., Bokharaei-Salim, F., Mirzaei, H., & Hamblin, M. R. (2020). Exosomal miRNAs: Novel players in viral infection. Epigenomics, 12(4), 353–370. 10.2217/epi-2019-0192

Nilsen, T. W. (2007). Mechanisms of microRNA-mediated gene regulation in animal cells. Trends in Genetics, 23(5), 243–249. 10.1016/j.tig.2007.02.011

O’Brien, J., Hayder, H., Zayed, Y., & Peng, C. (2018). Overview of MicroRNA Biogenesis, Mechanisms of Actions, and Circulation. Frontiers in Endocrinology, 9, 402. 10.3389/fendo.2018.00402

O’Donnell, K. A., Wentzel, E. A., Zeller, K. I., Dang, C. V., & Mendell, J. T. (2005). C-Myc-regulated microRNAs modulate E2F1 expression. Nature, 435(7043), Article 7043. 10.1038/nature03677

Olive, V., Bennett, M. J., Walker, J. C., Ma, C., Jiang, I., Cordon-Cardo, C., Li, Q.-J., Lowe, S. W., Hannon, G. J., & He, L. (2009). MiR-19 is a key oncogenic component of mir-17-92. Genes & Development, 23(24), 2839–2849. 10.1101/gad.1861409

Olive, V., Li, Q., & He, L. (2013). Mir-17-92, a polycistronic oncomir with pleiotropic functions. Immunological Reviews, 253(1), 158–166. 10.1111/imr.12054

Olive, V., Minella, A. C., & He, L. (2015). Outside the coding genome, mammalian microRNAs confer structural and functional complexity. Science Signaling, 8(368), re2. 10.1126/scisignal.2005813

Olive, V., Sabio, E., Bennett, M. J., De Jong, C. S., Biton, A., McGann, J. C., Greaney, S. K., Sodir, N. M., Zhou, A. Y., Balakrishnan, A., Foth, M., Luftig, M. A., Goga, A., Speed, T. P., Xuan, Z., Evan, G. I., Wan, Y., Minella, A. C., & He, L. (2013). A component of the mir-17-92 polycistronic oncomir promotes oncogene-dependent apoptosis. ELife, 2, e00822. 10.7554/eLife.00822

Omoto, S., & Fujii, Y. R. (2005). Regulation of human immunodeficiency virus 1 transcription by nef microRNA. Journal of General Virology, 86(3), 751–755. 10.1099/vir.0.80449-0

Omoto, S., Ito, M., Tsutsumi, Y., Ichikawa, Y., Okuyama, H., Brisibe, E. A., Saksena, N. K., & Fujii, Y. R. (2004). HIV-1 nef suppression by virally encoded microRNA. Retrovirology, 1(1), 44. 10.1186/1742-4690-1-44

Ota, A., Tagawa, H., Karnan, S., Tsuzuki, S., Karpas, A., Kira, S., Yoshida, Y., & Seto, M. (2004). Identification and Characterization of a Novel Gene, C13orf25, as a Target for 13q31-q32 Amplification in Malignant Lymphoma. Cancer Research, 64(9), 3087–3095. 10.1158/0008-5472.CAN-03-3773

Ouellet, D. L., Plante, I., Landry, P., Barat, C., Janelle, M.-E., Flamand, L., Tremblay, M. J., & Provost, P. (2008). Identification of functional microRNAs released through asymmetrical processing of HIV-1 TAR element. Nucleic Acids Research, 36(7), 2353–2365. 10.1093/nar/gkn076

Parisi F, Freer G, Mazzanti CM, Pistello M, Poli A. Mouse Mammary Tumor Virus (MMTV) and MMTV-like Viruses: An In-depth Look at a Controversial Issue. Viruses. 2022 May 6;14(5):977. doi: 10.3390/v14050977

Pfeffer, S., Sewer, A., Lagos-Quintana, M., Sheridan, R., Sander, C., Grässer, F. A., van Dyk, L. F., Ho, C. K., Shuman, S., Chien, M., Russo, J. J., Ju, J., Randall, G., Lindenbach, B. D., Rice, C. M., Simon, V., Ho, D. D., Zavolan, M., & Tuschl, T. (2005). Identification of microRNAs of the herpesvirus family. Nature Methods, 2(4), Article 4. 10.1038/nmeth746

Pfeffer, S., & Voinnet, O. (2006). Viruses, microRNAs and cancer. Oncogene, 25(46), Article 46. 10.1038/sj.onc.1209915

Pfeffer, S., Zavolan, M., Grässer, F. A., Chien, M., Russo, J. J., Ju, J., John, B., Enright, A. J., Marks, D., Sander, C., & Tuschl, T. (2004). Identification of Virus-Encoded MicroRNAs. Science, 304(5671), 734–736. 10.1126/science.1096781

Rani, V., & Sengar, R. S. (2022). Biogenesis and mechanisms of microRNA-mediated gene regulation. Biotechnology and Bioengineering, 119(3), 685–692. 10.1002/bit.28029

Rehmsmeier, M., Steffen, P., Hochsmann, M., & Giegerich, R. (2004). Fast and effective prediction of microRNA/target duplexes. RNA (New York, N.Y.), 10(10), 1507–1517. 10.1261/rna.5248604

Rennie, W., Kanoria, S., Liu, C., Carmack, C. S., Lu, J., & Ding, Y. (2019). Sfold Tools for MicroRNA Target Prediction. Methods in Molecular Biology (Clifton, N.J.), 1970, 31–42. 10.1007/978-1-4939-9207-2_3

Rennie, W., Kanoria, S., Liu, C., Mallick, B., Long, D., Wolenc, A., Carmack, C. S., Lu, J., & Ding, Y. (2016). STarMirDB: A database of microRNA binding sites. RNA Biology, 13(6), 554–560. 10.1080/15476286.2016.1182279

Rosewick, N., Momont, M., Durkin, K., Takeda, H., Caiment, F., Cleuter, Y., Vernin, C., Mortreux, F., Wattel, E., Burny, A., Georges, M., & Van den Broeke, A. (2013). Deep sequencing reveals abundant noncanonical retroviral microRNAs in B-cell leukemia/lymphoma. Proceedings of the National Academy of Sciences, 110(6), 2306–2311. 10.1073/pnas.1213842110

Ross, S. R. (2010). Mouse Mammary Tumor Virus Molecular Biology and Oncogenesis. Viruses, 2(9), Article 9. 10.3390/v2092000

Ross, S. R., Schmidt, J. W., Katz, E., Cappelli, L., Hultine, S., Gimotty, P., & Monroe, J. G. (2006). An immunoreceptor tyrosine activation motif in the mouse mammary tumor virus envelope protein plays a role in virus-induced mammary tumors. Journal of Virology, 80(18), 9000–9008. 10.1128/JVI.00788-06

Rouha, H., Thurner, C., & Mandl, C. W. (2010). Functional microRNA generated from a cytoplasmic RNA virus. Nucleic Acids Research, 38(22), 8328–8337. 10.1093/nar/gkq681

Rupaimoole, R., & Slack, F. J. (2017). MicroRNA therapeutics: Towards a new era for the management of cancer and other diseases. Nature Reviews Drug Discovery, 16(3), Article 3. 10.1038/nrd.2016.246

Salmons, B., Knedlitschek, G., Kennedy, N., Groner, B., & Ponta, H. (1986). The endogenous mouse mammary tumour virus locus Mtv-8 contains a defective envelope gene. Virus Research, 4(4), 377–389. 10.1016/0168-1702(86)90084-5

Shackleford, G. M., & Varmus, H. E. (1988a). Construction of a clonable, infectious, and tumorigenic mouse mammary tumor virus provirus and a derivative genetic vector. Proceedings of the National Academy of Sciences of the United States of America, 85(24), 9655–9659.

Shackleford, G. M., & Varmus, H. E. (1988b). Construction of a clonable, infectious, and tumorigenic mouse mammary tumor virus provirus and a derivative genetic vector. Proceedings of the National Academy of Sciences of the United States of America, 85(24), 9655–9659.

Shapiro, J. S., Varble, A., Pham, A. M., & tenOever, B. R. (2010). Noncanonical cytoplasmic processing of viral microRNAs. RNA, 16(11), 2068–2074. 10.1261/rna.2303610

Shi, B., Zhu, M., Liu, S., & Zhang, M. (2013). Highly ordered architecture of microRNA cluster. BioMed Research International, 2013, 463168. 10.1155/2013/463168

Slicing and dicing viruses: Antiviral RNA interference in mammals. (2019). The EMBO Journal, 38(8), e100941. 10.15252/embj.2018100941

Stenvang, J., Petri, A., Lindow, M., Obad, S., & Kauppinen, S. (2012). Inhibition of microRNA function by antimiR oligonucleotides. Silence, 3(1), 1. 10.1186/1758-907X-3-1

Stewart, A. F. R., & Chen, H.-H. (2022). Revisiting the MMTV Zoonotic Hypothesis to Account for Geographic Variation in Breast Cancer Incidence. Viruses, 14(3), 559. 10.3390/v14030559

Swanson, I., Jude, B. A., Zhang, A. R., Pucker, A., Smith, Z. E., & Golovkina, T. V. (2006). Sequences within the gag Gene of Mouse Mammary Tumor Virus Needed for Mammary Gland Cell Transformation. Journal of Virology, 80(7), 3215–3224. 10.1128/JVI.80.7.3215-3224.2006

Sylvestre, Y., De Guire, V., Querido, E., Mukhopadhyay, U. K., Bourdeau, V., Major, F., Ferbeyre, G., & Chartrand, P. (2007). An E2F/miR-20a autoregulatory feedback loop. The Journal of Biological Chemistry, 282(4), 2135–2143. 10.1074/jbc.M608939200

Tanzer, A., & Stadler, P. F. (2004). Molecular Evolution of a MicroRNA Cluster. Journal of Molecular Biology, 339(2), 327–335. 10.1016/j.jmb.2004.03.065

Thomson, D. W., Bracken, C. P., Szubert, J. M., & Goodall, G. J. (2013). On measuring miRNAs after transient transfection of mimics or antisense inhibitors. PloS One, 8(1), e55214. 10.1371/journal.pone.0055214

Toledo-Arana, A., Repoila, F., & Cossart, P. (2007). Small noncoding RNAs controlling pathogenesis. Current Opinion in Microbiology, 10(2), 182–188. 10.1016/j.mib.2007.03.004

Triboulet, R., Mari, B., Lin, Y.-L., Chable-Bessia, C., Bennasser, Y., Lebrigand, K., Cardinaud, B., Maurin, T., Barbry, P., Baillat, V., Reynes, J., Corbeau, P., Jeang, K.-T., & Benkirane, M. (2007). Suppression of MicroRNA-Silencing Pathway by HIV-1 during Virus Replication. Science, 315(5818), 1579–1582.

Umbach, J. L., Yen, H.-L., Poon, L. L. M., & Cullen, B. R. (2010). Influenza A Virus Expresses High Levels of an Unusual Class of Small Viral Leader RNAs in Infected Cells. MBio, 1(4), e00204–10. 10.1128/mBio.00204-10

Varble, A., Chua, M. A., Perez, J. T., Manicassamy, B., García-Sastre, A., & tenOever, B. R. (2010). Engineered RNA viral synthesis of microRNAs. Proceedings of the National Academy of Sciences, 107(25), 11519–11524. 10.1073/pnas.1003115107

Vilimova, M., & Pfeffer, S. (2023). Post-transcriptional regulation of polycistronic microRNAs. WIREs RNA, 14(2), e1749. 10.1002/wrna.1749

Wang, S., Liu, P., Yang, P., Zheng, J., & Zhao, D. (2017). Peripheral blood microRNAs expression is associated with infant respiratory syncytial virus infection. Oncotarget, 8(57), 96627–96635. 10.18632/oncotarget.19364

Wang, Z., Liu, M., Zhu, H., Zhang, W., He, S., Hu, C., Quan, L., Bai, J., & Xu, N. (2010). Suppression of p21 by c-Myc through members of miR-17 family at the post-transcriptional level. International Journal of Oncology, 37(5), 1315–1321.

Whisnant, A. W., Kehl, T., Bao, Q., Materniak, M., Kuzmak, J., Löchelt, M., & Cullen, B. R. (2014). Identification of Novel, Highly Expressed Retroviral MicroRNAs in Cells Infected by Bovine Foamy Virus. Journal of Virology, 88(9), 4679–4686. 10.1128/JVI.03587-13

Wilkins, C., Dishongh, R., Moore, S. C., Whitt, M. A., Chow, M., & Machaca, K. (2005). RNA interference is an antiviral defence mechanism in Caenorhabditis elegans. Nature, 436(7053), Article 7053. 10.1038/nature03957

Woods, K., Thomson, J. M., & Hammond, S. M. (2007). Direct regulation of an oncogenic micro-RNA cluster by E2F transcription factors. The Journal of Biological Chemistry, 282(4), 2130–2134. 10.1074/jbc.C600252200

Yang, H.-Y., Barbi, J., Wu, C.-Y., Zheng, Y., Vignali, P. D. A., Wu, X., Tao, J.-H., Park, B. V., Bandara, S., Novack, L., Ni, X., Yang, X., Chang, K.-Y., Wu, R.-C., Zhang, J., Yang, C.-W., Pardoll, D. M., Li, H., & Pan, F. (2016). MicroRNA-17 Modulates Regulatory T Cell Function by Targeting Co-regulators of the Foxp3 Transcription Factor. Immunity, 45(1), 83–93. 10.1016/j.immuni.2016.06.022

Zhang, Y., Fan, M., Geng, G., Liu, B., Huang, Z., Luo, H., Zhou, J., Guo, X., Cai, W., & Zhang, H. (2014). A novel HIV-1-encoded microRNA enhances its viral replication by targeting the TATA box region. Retrovirology, 11(1), 23. 10.1186/1742-4690-11-23

Zhang, Y., Yang, L., Wang, H., Zhang, G., & Sun, X. (2016). Respiratory syncytial virus non-structural protein 1 facilitates virus replication through miR-29a-mediated inhibition of interferon-α receptor. Biochemical and Biophysical Research Communications, 478(3), 1436–1441. 10.1016/j.bbrc.2016.08.142

Zhu, Y., Huang, Y., Jung, J. U., Lu, C., & Gao, S.-J. (2014). Viral miRNA targeting of bicistronic and polycistronic transcripts. Current Opinion in Virology, 7, 66–72. 10.1016/j.coviro.2014.04.004

